# Self-Organization of Sinusoidal Vessels in Pluripotent Stem Cell-derived Human Liver Bud Organoids

**DOI:** 10.1101/2024.07.02.601804

**Authors:** Norikazu Saiki, Yasunori Nio, Yosuke Yoneyama, Shuntaro Kawamura, Kentaro Iwasawa, Eri Kawakami, Kohei Araki, Junko Fukumura, Tsuyoshi Sakairi, Tamaki Kono, Rio Ohmura, Masaru Koido, Masaaki Funata, Wendy L. Thompson, Pamela Cruz-Encarnacion, Ya-Wen Chen, Takanori Takebe

## Abstract

The induction of tissue-specific vessels in *in vitro* living tissue systems remains challenging. Here, we directly differentiated human pluripotent stem cells into CD32b^+^ putative liver sinusoidal progenitors (iLSEP) by dictating developmental pathways. By devising an inverted multilayered air-liquid interface (IMALI) culture, hepatic endoderm, septum mesenchyme, arterial and sinusoidal quadruple progenitors self-organized to generate and sustain hepatocyte-like cells neighbored by divergent endothelial subsets composed of CD32b^low^CD31^high^, LYVE1^+^STAB1^+^CD32b^high^CD31^low^THBD^-^vWF^-^, and LYVE1^-^THBD^+^vWF^+^ cells. Wnt2 mediated sinusoidal-to-hepatic intercellular crosstalk potentiates hepatocyte differentiation and branched endothelial network formation. Intravital imaging revealed iLSEP developed fully patent human vessels with functional sinusoid-like features. Organoid-derived hepatocyte- and sinusoid-derived coagulation factors enabled correction of *in vitro* clotting time with Factor V, VIII, IX, and XI deficient patients’ plasma and rescued the severe bleeding phenotype in hemophilia A mice upon transplantation. Advanced organoid vascularization technology allows for interrogating key insights governing organ-specific vessel development, paving the way for coagulation disorder therapeutics.

## Main

Organoids are complex multicellular cultures that can be expanded and genetically manipulated, while possessing tissue microarchitecture and functionality^1–4^. Human organoid systems have facilitated unprecedented studies of human organ development and pathophysiology, holding therapeutic promise for patients with incurable diseases^5,6^. However, engineering the cellular complexity to more closely mimic individual organs’ native structures such as organ-specific vasculature remains a major challenge^7,8^.

During development and regeneration, organ-specialized blood vessels play essential roles in differentiating parenchymal lineages through direct and indirect interactions beyond establishing a passive circulatory conduit^9,10^. Previous studies highlighted the importance of VEGF-mediated angiocrine support for organogenesis and regeneration both in animal models and human models in liver^11–13^ and kidney^14,15^, suggesting the important roles of organ-specific endothelium in directing early organogenesis. For instance, in the developing liver, the endothelium produces soluble factors such as HGF, BMP, and Wnt ligands which are critical for hepatocyte growth and development^16^.

Liver-specific capillary plexus runs between radiating hepatic cords and is comprised of liver sinusoidal endothelial cells (LSECs)^17,18^. LSECs are morphologically unique among endothelial cells with minimal basement membrane and fenestrae that facilitate fluid exchange between the sinusoids and liver parenchyma^19^. Their location in the sinusoids gives LSEC’s access to systemic arterial and portal venous blood. LSECs are one of the most potent scavenger systems in the body, equipped with various scavenger receptors and strong endocytic capacity^20^. One significant clinical problem associated with LSEC involves hemophilia A, as LSECs are the main cell types to produce the coagulation Factor VIII (FVIII)^21,22^.

While previous studies have generated vascularized liver organoid cultures using human induced pluripotent stem cell (hiPSC)-derived endothelial cells (ECs), the resultant ECs, expressing arterial markers, failed to acquire LSEC characteristics^13,23,24^. Recently, several directed differentiation protocols generated progenitor cells from PSC-derived angioblasts^25–27^ that can engraft and mature LSEC characteristics following Monocrotaline-induced preconditioning^26,28^. However, the self-organization of these liver-specific ECs in human organoids represents a significant challenge.

Here, the prime goals of this study are to 1) develop the stepwise differentiation protocols for LSEC progenitors guided by longitudinal single-cell transcriptome during hepatic vascular development, 2) establish three-dimensional (3D) organoid culture conditions that allow self-organization of hepatic and endothelial subpopulations with key organ-specific functional features, 3) delineate intercellular signaling crosstalk that occurs during multicellular organoid development and 4) evaluate whether coagulation factors derived from these vascularized organoids can restore coagulation factor deficiencies both in animal and human models (**Fig. 1a**).

**Fig. 1:**
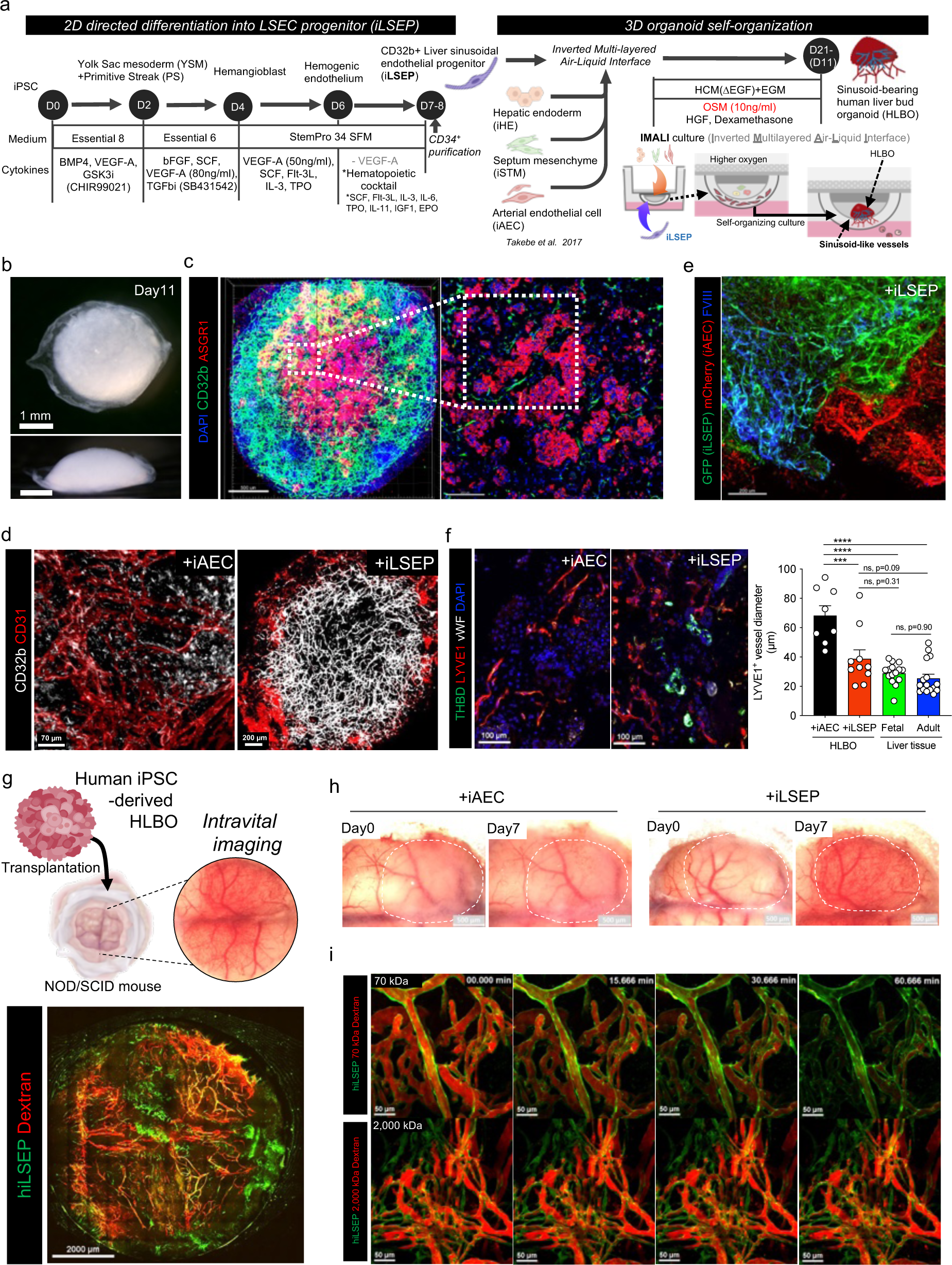
Self-organization of sinusoid-like endothelial network in iPSC-derived human liver bud organoids. a) Schematic overview of stepwise differentiation of iLSEC and generation of sinusoid- bearing HLBO by IMALI culture. b) Stereomicroscopic image of +iLSEP HLBO at day 11. Overall view from above (upper); side view (lower). c) 3D reconstructed image of HLBO with whole-mount immunofluorescent staining for CD32b and ASGR1 at day11 (left). Scale bar indicates 500 µm. Enlarged cross-sectional view of the white dashed square region within left panel (right). Scale bar indicates 200 µm. See **Supplementary Video 1** for detailed 3D structure. d) Whole-mount images of branched endothelial network in organoids with immunofluorescent staining for CD32b/CD31 at day 11. Right panel is enlarged image of the yellow dashed square region of left panel. e) Whole-mount image of HLBOs including GFP^+^ iLSEP and mCherry^+^ iAEC with immunofluorescent staining for FVIII at day 11. Scale bar indicates 200 µm. f) Cross-sectioning image of HLBOs with immunofluorescent staining for LYVE1/THBD/vWF at day 11. Right panels are quantification of LYVE1^+^ vessel diameter. The diameter of vessels with tubular structures was measured from the binarized images. Data represent the mean ± SEM (n=8,+ iAEC HLBO and +iLSEP HLBO; n=16; fetal and adult liver tissue; ****, p<0.0001; ***, p<0.001; ns, not significant; one-way ANOVA with Tukey’s multiple comparisons test). g) Schematic representation of HLBO transplantation under cranial window and intravital imaging (top). Intravital fluorescence microscopy imaging in the whole-window view of the +iLSEP HLBO transplanted mice (bottom). The organoid-derived human vessels were observed by GFP^+^ iLSEP. Blood flow was visualized using 2,000 kDa tetramethylrhodamine-dextran. h) Macroscopic image of transplanted HLBOs under cranial window. Dotted area indicates the transplanted HLBOs. Time-lapse imaging of 70 kDa or 2,000 kDa tetramethylrhodamine-dextran visualized- blood flow in GFP^+^ organoid-derived human vessels. See **Supplementary Video 2** for time-lapse movie.

## Results

### Directed differentiation of human PSC into liver sinusoid endothelial progenitor

During development, the branch of vitelline vein connecting between embryo and extraembryonic tissue (yolk sac) extends to septum transversum and subsequently undergoes remodeling of the primitive venous plexus. These vasculatures and endocardium of the sinus venosus from fetal embryonic heart collaboratively form part of the hepatic sinusoids^29–33^. We hypothesized that venous endothelial cells emanating from both embryonic and extraembryonic mesoderm can generate progenitors for early fetal liver ECs. To induce mesoderm progenitors including both primitive streak (PS) and yolk sac (YS) region, the 2D micropatterning differentiation method was applied^34,35^. Micropatterned colonies were then subjected to published hemangioblast differentiation protocols to gain yolk sac-like endothelial character (**Fig. 1a**)^36,37^. After day 6 of differentiation, the outer edge of colony expanded and began expressing CD31, an early vascular endothelial cell marker (**Extended Data Fig. 1a**, b). In the spreading CD31^+^ cell area located between the outer edge and the center, hemogenic endothelial cells expressing CD32b, a subfamily of Fc gamma receptor 2 that is expressed in the YS endothelial region^38^, were emerging (**Extended Data Fig. 1c**). A high concentration of VEGF-A is prohibitive for specification into venous-like endothelial cells^26,39^. We next attempted to induce venous LSEC progenitors by exposing day 6 hemogenic endothelium to a hematopoietic cytokine cocktail without VEGF-A. After two days-treatment with the hematopoietic cytokine cocktail, flow cytometry analysis showed that the fraction of sorted day 8 CD34^+^ cells co-expressed CD31 and CD32 reached 88 ± 0.7% purity (**Extended Data Fig. 1d**). To further define the CD34^+^CD31^+^CD32^+^ cell population, we performed bulk RNA seq analysis and compared gene expression levels of our iPSC-derived putative liver sinusoidal endothelial progenitors (iLSEP) to primary LSECs, iPSCs, and the induced septum transversum mesenchyme (iSTM) and conventional ECs used in our previously reported liver bud organoid^24^. iLSEPs widely expressed more markers such as venous EC (*NR2F2*, *PROX1*, and *APLNR*), endocardium (*CDH11*, *CTHRC1*, and *NPR3*), yolk sac mesoderm (*SPP1*, *TPM2*, *TTR*, *FOXF1*, and *HAND1*), and LSEC markers (*STAB2*, *LYVE1*, and *FCGR2B*) (**Extended Data Fig. 1e**). In contrast, conventional ECs differentiated using a high VEGF-A concentration (100 ng /mL) are more highly enriched for arterial genes than those expressed in LSECs and iLSEPs such as CD32, hereafter defined as induced arterial endothelial cells (iAECs) (**Extended Data Fig. 1d**, e). In the venous angioblasts (VAs) differentiated by the embryoid body (EB)-based protocol^26^, expression of CD32 and CD31 was heterogeneous (**Extended Data Fig. 1f**). Thus, we established a stepwise protocol for 2D differentiation of iLSEP as a putative progenitor for LSEC, from micropatterned colonies.

Next, to capture the key transcriptional changes that occur during the acquisition of LSEC fate^40,41^ in hepatic vascular development, we integrated single-cell gene expression profiles of the adult liver, fetal liver, yolk sac, and fetal heart and extracted a dataset containing endocardium, yolk sac ECs, early, mid, and late fetal liver ECs (divided by post-conception week (PCW)), and adult liver midzonal and peri-central (zone 2 and 3) sinusoidal ECs, classified according to the annotations defined by previous reports^40^. A trajectory map on UMAP recapitulated the developmental time series of LSECs from endocardium and yolk sac EC (**Extended Data Fig. 2a**). Pseudo-time reconstructed profiling confirmed a decrease in YS and endocardium markers and an increase in LSEC markers according to the developmental stages (**Extended Data Fig. 2b**, c and **Supplementary Table 1, 2**). Pathway enrichment analysis on the gene modules during fetal LSECs differentiation showed the early activation of vascular proliferation signaling pathways such as pathways associated with VEGF-A^13,42^, Gastrin^43^ and Oncostatin M (OSM)^44^ before complement and coagulation pathways are upregulated from the later developmental stage to the adult stage (**Extended Data Fig. 2d**, e, and **Supplementary Table 3, 4**). To test for the role of OSM in sinusoidal fate induction, we purified iLSEPs by selecting the CD34^+^ fraction and then treated with or without OSM (**Extended Data Fig. 2f**). At day14, when OSM was added to iLSEPs, the purity of CD31^+^, CD32^+^, and LYVE1^+^ sinusoidal endothelial-like cell populations increased in an OSM concentration-dependent manner, reaching 90.1 ± 1.2% (**Extended Data Fig. 2g**, h). Consistent with the integrated single-cell gene expression data (**Extended Data Fig. 2i**), OSM-treated iLSEP converted into a highly homogenous CD73^+^CXCR4^-^ population (93.2 ± 1.3%, **Extended Data** Fig 2j). Immunocytochemical staining revealed that CD32b^+^ cells (both iLSEP and iLSEC) expressed higher levels of FVIII than CD31^+^CD32b^-^ cells (iAEC), and CD32b and FVIII expression were enhanced upon differentiation of iLSEP to iLSEC (**Extended Data Fig. 2k**). The differentiation protocol from VA to LSEC-like cells induced high purity LYVE1^+^CD32^+^ cells (VA-LSEC)^26^ as in our iLSECs (**Extended Data Fig. 2l**), but the protein expression of FVIII was significantly weak (**Extended Data Fig. 2m**). Secretion levels of FVIII were significantly increased in iLSEC compared to VA-LSEC, iAEC, and iLSEP (**Extended Data Fig. 2n**). Fenestrae or cellular membrane micropores are typical characteristics of LSECs, that connect the lumen of sinusoids with the space of Disse^19^. Scanning electron microscopy (SEM) indicated the presence of fenestrae in iLSEC that is absent in iAEC (**Extended Data Fig. 2o**). These results suggest that iLSEPs possess differentiation capacity to LSEC-like cells upon OSM signal activation as seen in fetal LSEC development.

### Controlled self-assembly into vascularized liver organoids

Exploration of multi-cellular self-organization necessitates 3D culture techniques that allow studying anatomical boundaries in organoid systems, while maintaining the necessary oxygenation and nutrient conditions for sustaining multicellular health. To accomplish this with liver bud organoids^24^ and iLSEC progenitors, we developed an inverted multilayered air-liquid interface 3D (IMALI) culture using porous membrane filters combined with a multilayering technique (**Fig. 1a**)^45,46^. IMALI system recapitulates the anatomical relationship between the primitive vascular plexus and liver bud during early development. While sufficient oxygenation is provided from the surface, nutrients are supplied from the bottom. First, we embedded iPSC-derived hepatic endoderm (iHE), iSTM, and iAEC in a first layer adjacent to the membrane, followed by an additional layering of either iLSEPs or iAECs. Over the next 5 days, the cells condensed into spheroids to self-organize into human liver bud organoids (HLBO). The HLBO culture with iLSEP displayed a more tightly packed morphology than the iAEC alone cultures, forming a dense dome-like structure, approximately 3.0 mm × 3.0 mm × 1.5 mm in size, around day 11 (**Fig. 1b** and **Extended Data Fig. 3a**). Whole mount immunofluorescent staining demonstrated that iLSEPs, but barely iAEC, formed a branched 3D endothelial network inside HLBOs containing both CD32b^+^ CD31^+^ cells and CD32b^-^ CD31^+^ cells, neighbored by ASGR1+ hepatocyte clusters (**Fig. 1c, d,** and **Supplementary Video 1**). Higher magnification images confirmed that iLSEP-derived CD32b^+^ cells express the LSEC-specific FVIII that was missing in iAEC alone culture (**Fig. 1e** and **Extended Data Fig. 3b**, c). Flow cytometry analysis showed that 90.5% of iLSEP-derived CD34^+^ endothelial cells in HLBO expressed CD32 (**Extended Data Fig. 3d**). In detail, there were three endothelial cell populations, CD32^high^CD31^low^, CD32^low^CD31^low^, and CD31^high^ found in our iLSEP in HLBO cultures, and of these populations, CD32^high^CD31^low^ cells and some CD32^low^CD31^low^ cells expressed CD36, a scavenger receptor expressed on LSECs^47,48^ (**Extended Data Fig. 3e**). Immunofluorescence analysis further revealed the presence of both central venous-like EC subset in the HLBO (THBD^+^vWF^+^LYVE1^-^)^49^ and sinusoidal-like EC (LYVE1^+^STAB1^+^CD32b^+^THBD^-^vWF^-^) which form a branched network with similar tubular diameter to those in fetal and adult liver tissue (**Fig. 1f** and **Extended Data Fig. 3f**). The microstructure of LYVE1-expressing sinusoidal EC in HLBO, visualized by transmission immunoelectron microscopy, showed a porous and fenestrae-like structure^50^ (**Extended Data Fig. 3g**).

To examine *in vivo* vascular network reconstitution capability, HLBOs were implanted into cranial window of immunodeficiency mice for the subsequent optical access^51^. Intravital imaging showed that the +iLSEP HLBO developed patent and fully perfused blood vessels within 6 days post-transplantation and further remodeled to form functional vasculatures (**Fig. 1g, h** and **Extended Data Fig. 3h**). It has been shown that a 70 kDa molecular weight, but not 2,000 kDa, dextran is permeable in the hepatic sinusoidal vessels^52^. Thus, the permeability of organoid-derived vessels was tested by injection of two different molecular weight fluorescent dextran. In +iLSEP HLBO transplant, 2,000 kDa dextran was fully constrained within the reconstructed blood vessels while 70 kDa dextran was permeated out of the vessel (**Fig. 1i** and **Supplementary Video 2**). *In vivo* injection of acetylated-low density lipoprotein (AcLDL) dye confirmed the uptake in human endothelial cells that are specific to +iLSEP HLBO (**Extended Data Fig. 3i**). These results suggest formation of functional sinusoid-like vascular network in HLBO transplant.

### Organ-specific cell signatures and diversity displayed by single cell transcriptomics

To uncover cellular characteristics during organoid development at transcriptomic level, time-series single cell RNA-seq (scRNA-seq) data were acquired. First, HLBOs at three culture time points were integrated and UMAP embedding and hierarchical clustering were performed. For the identified populations, six cell types and 32 subtypes (three of which were unidentified) were assigned based on known markers (**Fig. 2a**, **Extended Data Fig. 4a** and **Supplementary Table 5**). At day 3, we found immature and proliferative hepatoblasts, along with several types of endothelial cells, proliferative mesenchymal cells, and other cardiovascular mesoderm progenitor cells (**Extended Data Fig. 4b**, c). At day 11, these progenitor subsets further differentiated into hepatocytes, cholangiocytes, endothelial cells, and stellate cells, with common marker gene expression found in fetal and adult liver cells (**Extended Data Fig. 4b**-e).

**Fig. 2:**
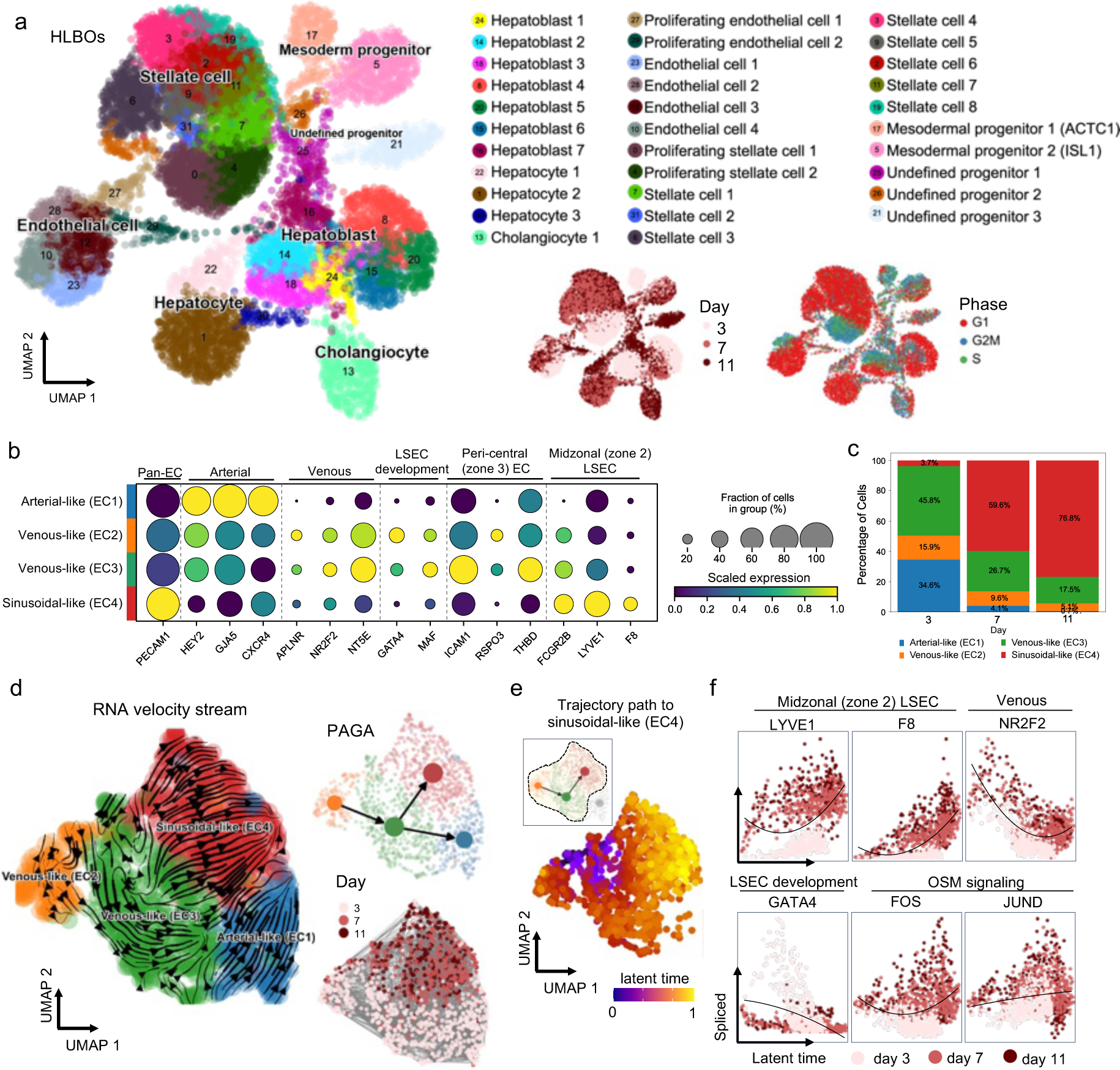
Profiling of endothelial trajectory and subpopulation in HLBO by longitudinal scRNA-seq. a) UMAPs for integrated +iLSEP HLBO dataset including day 3, 7, and 11 colored by louvain clustering group with cell type annotation, culture days, or cell cycle phase. b) The dot plot representing gene expression of endothelial marker genes that changed among each subtype group. The size of dots indicates the relative gene expression in percent for each group. The color represents the average expression level for the indicated gene. c) Proportion of endothelial subtype in each culture time series. d) Diffusion stream estimated by RNA velocity analysis (left) and partition-based graph abstraction (PAGA) graph (top right) overlaid on the UMAP colored by endothelial subtype. UMAP colored by culture time series (bottom right) e) UMAP showing extracted subpopulations on the trajectory path to sinusoidal-like endothelial cell (EC4) (upper left) and an inferred latent time. Scatter plots showing latent time versus spliced gene expression of endothelial markers. Black line on scatter plot is linear regression fitted line to represent trends of change.

Next, we constructed a dataset of 91,700 cells using a curated collection of 11 publicly available human PSC-derived liver organoid datasets^2,13,53–58^ to benchmark our HLBO model against existing models using neighborhood graph correlation mapping and spectrum comparison^59,60^ (**Extended Data Fig. 5a** and **Supplementary Table 6**). With respect to the status of hepatocytes and cholangiocytes, the +iLSEP HLBO showed a higher correlation with primary liver tissue at the later culture days (**Extended Data Fig. 5b**-d), albeit minimal difference in stellate cells (**Extended Data Fig. 5b**, e). For endothelial cells, the profiles of the +iLSEP HLBO at day 11 showed the highest correlation with primary LSECs, with spearman correlation values comparable for both adult and fetal (**Extended Data Fig. 5b**, f). To investigate iLSEP-derivatives, endothelial cell populations were extracted to analyze zone 2 and 3 LSEC markers. Only HLBO develops cells that satisfied all four LSEC marker expressions (**Extended Data Fig. 6a**). Extracted population were further defined into four clusters based on marker expression: arterial-like (EC1), venous-like (EC2, EC3) and sinusoidal-like endothelial cells (EC4) (**Fig. 2b** and **Supplementary Table 7**). Compositional analysis indicates relative expansion of CD32^+^F8^+^LYVE1^+^ sinusoidal endothelial cells over time, relative to LYVE1^-^THBD^+^ICAM1^+^ venous and CD31^+^CD32b^-^ arterial subtypes (**Fig. 2c**). The endothelial diversity was not apparent in the other organoid platforms (**Extended Data Fig. 6b**).

RNA velocity stream tracing subtype composition trajectories suggested a differentiation trajectory from venous to sinusoidal transition, in agreement with our differentiation hypothesis (**Fig. 2d**). Kinetic plot with an inferred latent time revealed that LYVE1^+^F8^+^ sinusoidal-like EC (EC4) was distinct from NR2F2^+^ venous-like ECs (EC2 and EC3) (**Fig. 2e**). Comparing the expression dynamics of each gene, sinusoidal markers were synchronized with the expression of transcription factors downstream of oncostatin M signaling, and GATA4, a key regulator of LSEC specification in liver development^61,62^, was preferentially expressed at day 3 (**Fig. 2f**). In arterial-like EC (EC1), arterial markers expression decreased along latent time, while NR2F2 expression increased (**Extended Data Fig. 6c**). Collectively, the scRNA-seq analysis elucidated *in vivo* relevant unique molecular profiles of multiple endothelial subsets in HLBO relative to the other organoid technology.

### Hepatic functionalization through the inclusion of iLSEP

In our co-culture system, we noted that the level of AFP mRNA in +iLSEP HLBO was lower than +iAEC HLBO (**Extended Data Fig. 7a**). In contrast, ALB gene expression and protein secretion were higher in the +iLSEP HLBO (**Extended Data Fig. 7a**, b). Hematoxylin and eosin staining of transverse sections of HLBO showed a mostly uniform distribution of hepatocyte-like clusters in +iLSEP HLBO (**Fig. 3a**). Further immunostaining revealed hepatocytes with differential expression levels of metabolic enzymes such as glutamine synthetase (GS), and glutaminase 2 (GLS2), were segregated (**Fig. 3b**). The percentage area of expression being GS, GLS2 double positive: 44.0 ± 3.0%, GS single positive: 44.4% ± 2.4%, and GLS2 single positive: 11.5% ± 2.2%, respectively, a different composition from the distribution biased toward GLS2 single (69.0% ± 4.2%), as in +iAEC (**Fig. 3c**). GS expressing cells located adjacent to GFP-tagged iLSEP-derived LYVE1^+^ endothelial cell within +iLSEP HLBO, while not seen in +iAEC HLBO, no endothelial cell HLBO (-EC), and non-layered air-liquid interface (ALI) culture condition (Non-layered ALI) (**Fig. 3c** and **Extended Data Fig. 7c**, d). Whole-mount immunostaining of HLBOs displayed that ASGR1+ hepatocytes, surrounded by CD32b^+^ sinusoidal ECs, expressed more GS in the +iLSEP HLBO than in +iAEC HLBO (**Fig. 3d, e** and **Extended Data Fig. 7e**). In parallel, Wnt downstream genes were also more active in the +iLSEP HLBO (**Extended Data Fig. 7f**). Consequently, metabolic activity of glutamine synthesis and ammonia uptake were thereby upregulated (**Fig.3f** and **Extended Data Fig. 7g**, h). Drug metabolism enzymes CYP3A4 and CYP1A2 typically activated in the peri-central region were efficiently induced by rifampicin and omeprazole, respectively (**Fig. 3g** and **Extended Data Fig. 7i**, j). Gene set enrichment analysis comparing hepatocyte of published human organoid datasets indicated that hepatic function associated genes, such as amino acid metabolism, cholesterol synthesis, and the coagulation and complement systems, were upregulated in the +iLSEP HLBO (**Extended Data Fig. 7k**). Consistently, complement proteins were highly excreted in the culture supernatant of the +iLSEP HLBO compared to the +iAEC and -EC HLBO (**Fig. 3h**). These results indicate that the inclusion of iLSEP promotes the emergence of hepatocytes with metabolic and protein secretion function in liver organoids.

**Fig. 3:**
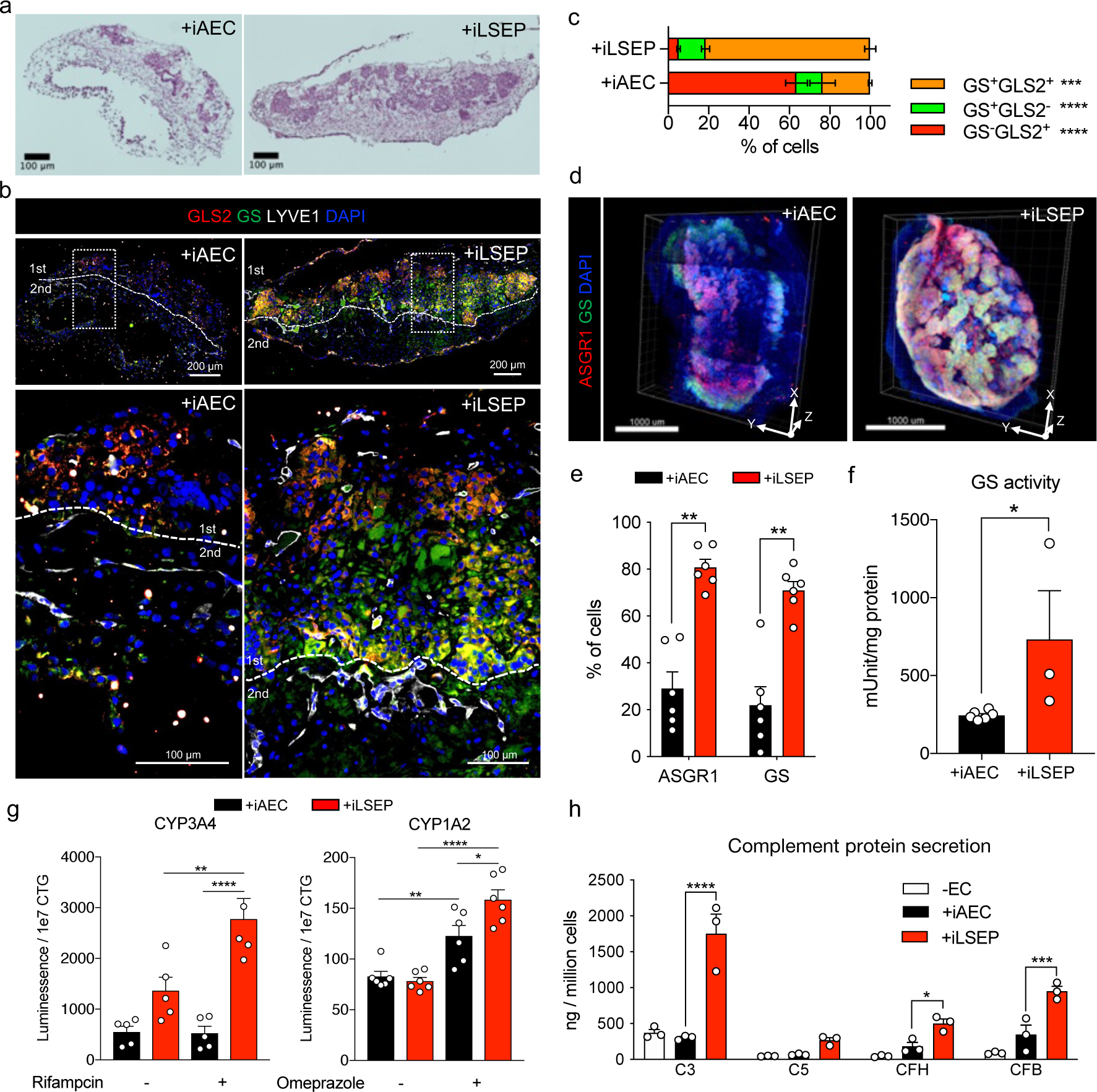
iLSEP promotes the maturation and functionalization of hepatocytes in HLBO a) Cross-sectioning image of HLBOs with H&E staining. b) Distribution of hepatocytes with different metabolic enzymes identified by immunofluorescent staining for GS (peri-central) and GLS2 (pan-lobular) in HLBOs. Lower panels are enlarged images of the white dashed square region. The yellow dashed lines indicate the boundaries of the gel layer in organoid generation as illustrated in Fig. 1a. c) Quantification of the percentage of cells expressing GS or GLS2 alone or co-expressing GS and GLS2. Segmentation and counting were normalized by counterstaining (DAPI). Data represent the mean ± SEM (n=4-8; ***, p<0.001; ****, p<0.0001; multiple t-test with Holm-Sidak correction) d) Whole-mount 3D reconstructed image of HLBOs with immunofluorescent staining for GS and ASGR1. Resolution is 2.8 µm × 2.8 µm × 8.2 µm per voxel. e) Quantification of the percentage of cells expressing GS or ASGR1 in 3D reconstructed images. Total cell numbers were counted by counterstaining (DAPI). Data represent the mean ± SEM (n=6; **, p<0.01; Mann-Whitney’s U-test). f) Enzymatic activity of GS in HLBOs protein extracts at day 11. Data represent the mean ± SEM (n=3-6; *, p<0.05; Mann-Whitney’s U-test) g) Measurement of CYP3A4 and CYP1A2 enzyme activities of organoids, with activation by rifampicin for CYP3A4 or omeprazole for CYP1A2. Data represent the mean ± SEM (n=5, CYP3A4; n=6, CYP1A2; *, p<0.05; ***, p<0.001; ****, p<0.0001; two-way ANOVA with Tukey’s multiple comparisons test) Measurement of complement proteins in organoid supernatants at day11. Data represent the mean ± SEM (n=3; *, p<0.05; ***, p<0.001; ****, p<0.0001; one-way ANOVA with Dunnett’s multiple comparisons test compared to +iLSEP HLBO).

### Endothelial-to-hepatic intercellular signaling supports HLBO functionalization

To further understand intercellular mechanisms during organoid formation, we analyzed the transcriptional profile of angiocrine factors in HLBO by RNA-seq. Hierarchical clustering exposed unique gene clusters containing angiocrine factors such as WNT ligands, *ANGPT1*, and *HGF* that were highly expressed in the iLSEPs (**Fig. 4a**). We inferred key intercellular signaling using ligand-receptor analysis. By counting the number of signaling pairs, iLSEP were found to be an epicenter for angiocrine communications (**Fig. 4b**). For example, WNT ligands that were upregulated in iLSEP were most predominantly received by iHE along with iSTM synergizing activation of WNT signaling (**Fig. 4c**). Similar interactions were found in in late-stage human fetal liver, where the WNT2-FZD5/6 ligand-receptor pair was significantly enriched (**Extended Data Fig. 8a**, b). This WNT2-mediated communication signature was consistent between iLSEP and iHE within the HLBO (**Fig. 4d**). Further exploration of the responder signals showed that the iHE produced VEGF-A, F2, PLG, GDF15, PROC, and EPO that can interact with receptors on iLSEPs. These have been shown to promote fetal liver and organoid maturation through multi-lineage interaction (**Extended Data Fig. 8c**-e)^13^.

**Fig. 4:**
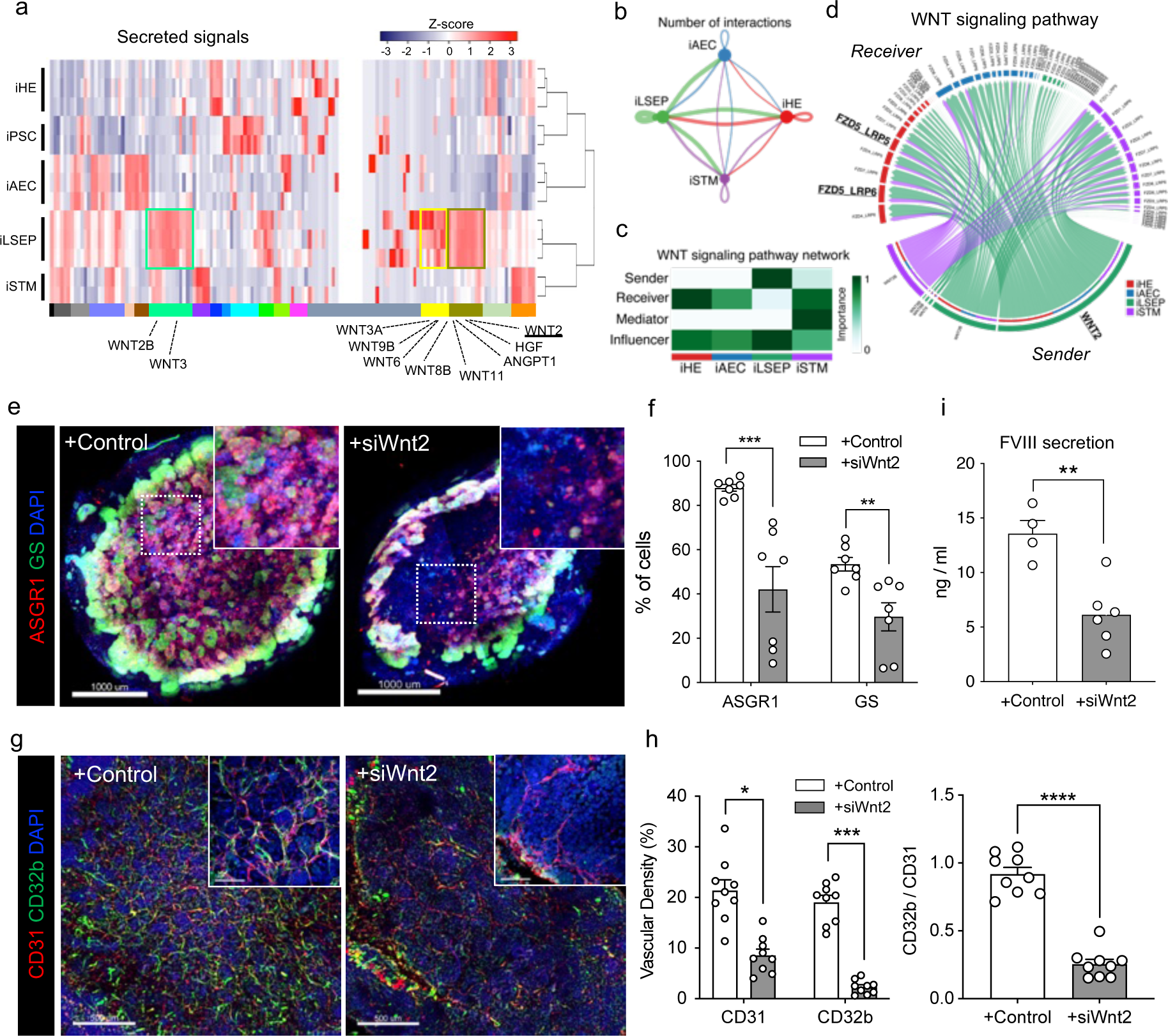
Endothelial to hepatic intercellular signal communication supports HLBO functionalization. a) The heatmap showing cytokines and growth factors associated gene set derived from GO class: ‘GO_SIGNALING_RECEPTOR_BINDING’. b) The chord diagram showing the intercellular communication network within the HLBO and the number of significant ligand-receptor pairs by the thickness of their edges (p<0.05). c) The heatmap visualizing the relative importance of each cell type of HLBO based on the centrality score of WNT signaling pathway. d) The chord diagram describing significant ligand-receptor pairs involved WNT signaling within HLBO (p<0.05). The ligand and receptor names that overlap with intercellular communication in fetal liver are highlighted. e) Whole-mount image of siRNA-treated HLBOs with immunofluorescent staining for GS and ASGR1. Scale bar indicates 500 µm. f) Quantification of the percentage of cells expressing GS or ASGR1 in 3D reconstructed images. Total cell numbers were counted by counterstaining (DAPI). Data represent the mean ± SEM (n=6; **, p<0.01; ***, p<0.001; Mann-Whitney’s U-test). g) Whole-mount image of siRNA-treated HLBOs with immunofluorescent staining for CD32b and CD31. h) Quantification of vascular density (% area of vessels) and ratio of CD32b^+^ / CD31^+^ area. Data represent the mean ± SEM (n=9; *, p<0.05; **, p<0.01; ***, p<0.001; ****, p<0.0001; one-way ANOVA with Tukey’s multiple comparisons test). i) Measurements of FVIII in HLBO supernatants. Data represent the mean ± SEM (n=5-12; *, p<0.05; ***, p<0.001; one-way ANOVA with Tukey’s multiple comparisons test).

WNT2 is known to be a major central venous SEC-derived activator of the beta- catenin signal controlling cell proliferation and metabolic zonation^12,16,63,64^. In 2D cultures, the knockdown of Wnt2 by siRNA had minimal influence on iLSECs characteristics (**Extended Data Fig. 9a**-c). We then performed IMALI culture-specific knockdown of Wnt2 by mixing lipid nanoparticles (LNPs)-siRNA complex only in the gel layer containing GFP-tagged iLSEPs (**Extended Data Fig. 9d**-f). Reduction of Wnt2 signaling resulted in a decrease in the number of ASGR1- and GS-expressing hepatocytes (**Fig. 4e, f** and **Extended Data Fig. 9g**). In parallel, iLSEP-specific Wnt2 knockdown compromised VEGF-A secretion from the HLBO (**Extended Data Fig. 9h**). Immunostaining of vascular endothelial markers showed a decreased branched endothelial network density and a lower percentage of CD32b^+^ LSECs among CD31^+^ pan- endothelial cells in the siWnt2-treated organoids (**Fig. 4g, h**). Similarly, LYVE1+ endothelial fractions were significantly reduced, while STAB1 expression remained unaffected by siWnt2 treatment (**Extended Data Fig. 9i**-k). LSEC-derived coagulation factor, FVIII^21^ was also decreased by Wnt2 knockdown (**Fig. 4i**). In sum, the interaction by Wnt2 signaling-mediated reciprocal crosstalk is essential for hepatocyte and endothelial differentiation.

### *In vitro* and *in vivo* coagulation factor functions in sinusoid-bearing HLBOs

One vital role of the liver is to synthesize and secrete various coagulation proteins, initiated even at fetal stages in the developing liver^65^. Given high levels of the diverse coagulation gene expression (**Fig. 5a**), we set out to study the functional coagulator activity of secreted proteins from our HLBOs. To determine the dynamics of functional FVIII production, we measured FVIII which was secreted into the media from the organoids over time. The +iLSEP HLBO had a significantly higher level of FVIII secretion (3 to 5-fold increase) than the +iAEC or -EC HLBOs, and FVIII production increased through 30 days of culture in the +iLSEP HLBO (**Fig. 5b**).

**Fig. 5:**
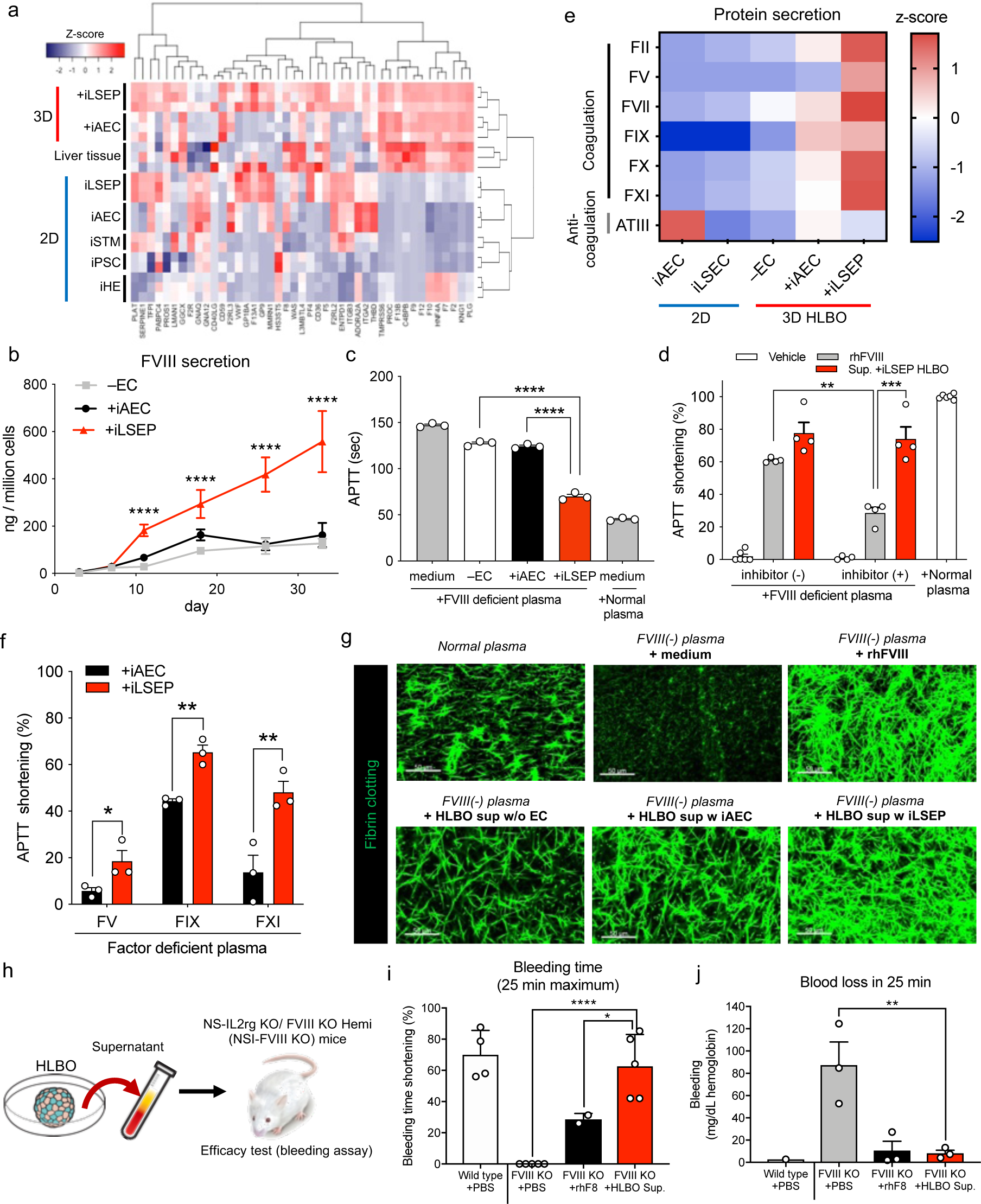
*in vitro* and *in vivo* coagulation factor functions in sinusoid-bearing HLBOs. a) The heatmap showing coagulation-associated gene set. b) Time series measurements of FVIII in organoid supernatants. Data represent the mean ± SEM (n=6; ****, p<0.0001; one-way ANOVA with Dunnett’s multiple comparisons test compared to +iLSEP HLBO). c) Activated partial thromboplastin time (APTT) reflecting FVIII activity of organoid supernatants at day 11 in FVIII deficient plasma compared to non-cultured media and normal reference plasma. Data represent the mean ± SEM (n=3; ****, p<0.0001; one-way ANOVA with Dunnett’s multiple comparisons test compared to +iLSEP HLBO). d) APTT shortening rate of +iLSEP HLBO supernatants at day 11 and recombinant human FVIII (rhFVIII) diluted to the same concentration of FVIII with inhibitor (+) or (-) FVIII deficient plasma relative to non-cultured media. Data represent the mean ± SEM (n=4; **, p<0.01; ***, p<0.001; one-way ANOVA with Sidak’s multiple comparisons test among rhFVIII and Sup. +iLSEP HLBO groups). e) Coagulation-associated factor profile in 2D-EC and 3D-organoid supernatants at day 11. Color values indicate z-score based on log2 transformed concentration (n=3). f) APTT shortening rate of HLBO supernatants at day 11 in factor (FV, FIX, and FXI) deficient plasma relative to non-cultured media. Data represent the mean ± SEM (n=3; *, p<0.05; **, p<0.01; multiple t-test with controlling FDR). g) Fibrin clotting images using Alexa Fluor 488-conjugated fibrinogen. h) Schematic representation of *in vivo* efficacy test with an injection of the culture supernatant of HLBO at day 11. See **Supplementary Video 3** for the bleeding assay. i) Shortening of time until bleeding stopped relative to maximum (25 min). Data represent the mean ± SEM (n=4-5; **, p<0.01; one-way ANOVA with Dunnett’s multiple comparisons test compared to FVIII KO mice with HLBO supernatant injection). Total blood loss during 25 min bleeding evaluated by hemoglobin concentration. Data represent the mean ± SEM. Dots that are not displayed indicate that the data was not detected (n=3; **, p<0.01; one-way ANOVA with Dunnett’s multiple comparisons test compared to FVIII KO mice with HLBO supernatant injection).

Hemophilia A is characterized by a reduced or absent FVIII activity, wherein the standard treatment is the repeated intravenous infusion of recombinant human FVIII (rhFVIII) protein^66^. To determine if HLBO-derived coagulation factors allow for the correction of Hemophilia A pathology, we combined patient-derived FVIII deficient plasma with the supernatant from HLBO with either iLSEP, iAEC or no EC and analyzed clot formation. An activated partial thromboplastin time (APTT) assay and demonstrated a 4-fold higher shortening of APTT relative to the +iAEC HLBO at day 11 and this effect lasted at least for 35 days (**Fig. 5b, c**). Compared to the reported LSEC progenitor VAs, the incorporation of iLSEPs in HLBO (**Extended Data Fig. 10a**) demonstrated the better performance in FVIII secretion and APTT restoration (**Extended Data Fig. 10b**-d). Interestingly, when the APTT assay was performed using Hemophilia A patient plasma carrying a FVIII inhibitor, the coagulation activity was reduced by about 50% in the recombinant FVIII group, while its efficacy remains almost intact in the +iLSEP HLBO supernatant group (**Fig. 5d**). We also found that the +iLSEP HLBO had the highest amounts of arrays of key coagulation factors (FII, FV, FVII, FIX, FX, and FXI) (**Fig. 5e**). With patients-plasma deficient for FV, FIX and FXI, the APTT shortening in +iLSEP HLBO are remarkably higher than in +iAEC (**Fig. 5f**). The +iLSEP HLBO supernatant restored fibrin clot formation of FVIII deficient plasma at comparable levels to rhFVIII (**Fig. 5g**).

Next, we generated immunodeficient NS-IL2rg KO/ FVIII KO Hemi (NSI-FVIII KO) mice to study the *in vivo* efficacy (**Extended Data Fig. 10e**). HLBO-derived secretome were administered into hemophilia A mice (**Fig. 5h**). Ultra-filtrated HLBO culture supernatant concentrates were injected intravenously into NSI-FVIII KO mice, and subsequently, a bleeding assay was performed by cutting each mouse’s tail-tip to confirm continuous bleeding and/or clotting. In control NSI-FVIII KO mice, tail-tip bleeding persisted for over 25 min (**Fig. 5i**). In contrast, in the mice infused with HLBO-derived proteins, bleeding ceased quickly at a similar timeframe as the current standard therapy using rhFVIII (**Fig. 5i, j** and **Supplementary Video 3**). Thus, +iLSEP HLBO produces arrays of functional coagulation proteins that can restore clot function in multiple hemorrhagic patient-derived serum assays and Hemophilia A model.

### *In vivo* hemophilia A correction by orthotopic transplantation of sinusoid-bearing HLBOs

To examine continued FVIII function *in vivo*, we transplanted HLBO and assessed coagulation function in an immunodeficient hemophilia A mouse model. Orthotopically transplanted HLBO successfully engrafted and reconstituted human CD32b^+^ vasculatures with functional FVIII secretion (**Fig. 6a-c**). The transplanted organoids expressed human FVIII in CD32b^+^LYVE1^+^ Ku80^+^ human endothelial cells (**Fig. 6b** and **Extended Data Fig. 10f**). Human FVIII protein secreted from HLBO was detected and significantly increased over time (**Fig. 6c**). Among various coagulation factors (**Fig. 5e**), FVIII was the most prominent factor that showed a significant increase in circulation in the plasma of transplanted mice (**Extended Data Fig. 10g**). To test *in vivo* efficacy, we performed a bleeding assay after transplantation. In all mice with successfully engrafted organoids, bleeding time and blood loss were clearly reduced at 4 and 8 weeks (**Fig. 6d-f** and **Supplementary Video 4**). Functionally active FVIII production continued for at least 5 months, measured by the APTT assay (**Fig. 6c, g**). The iLSEP incorporation into HLBOs was essential to exert therapeutic potential of FVIII production (**Extended Data Fig. 10h**). Transplantation of iLSECs alone failed to ameliorate bleeding phenotype (**Extended Data Fig. 10i**, j). Collectively, our vascularized organoid transplantation approach successfully restores the hemophilia A phenotype.

**Fig. 6:**
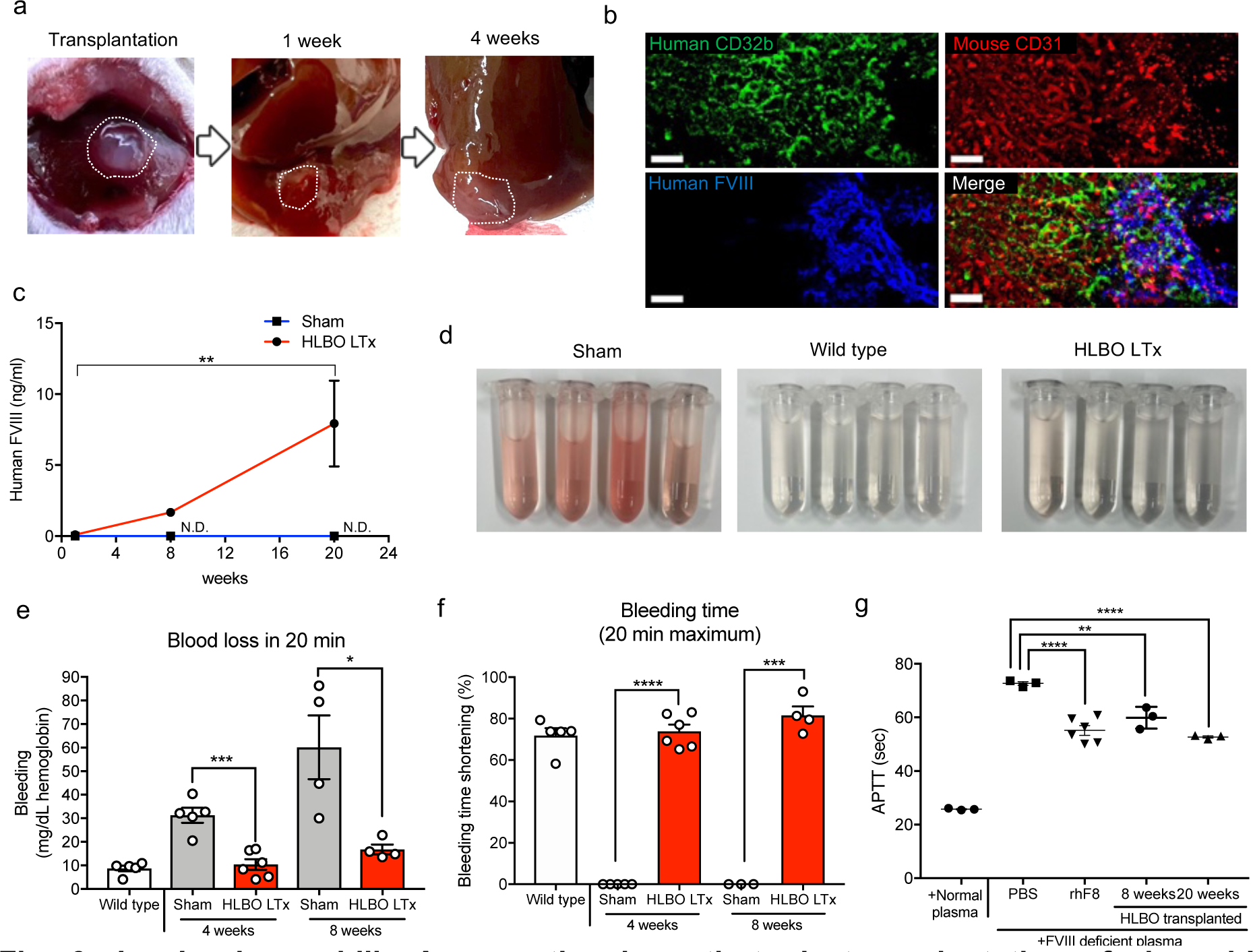
*in vivo* hemophilia A correction by orthotopic transplantation of sinusoid- bearing HLBOs. a) Morphological representation of transplanted HLBO into liver. b) Vascular network connection and overlapping between mouse-CD31^+^ host vessels and human-CD32b^+^FVIII^+^ vessels visualized by whole-mount image of with immunofluorescent staining. Scale bar indicates 50 µm. c) Time-series measurement of HLBO derived FVIII circulating in mouse blood 1, 8, and 20 weeks after liver transplantation (LTx) (n=3-5, transplanted; n=3, sham; **, p<0.01; two-way ANOVA with Tukey’s multiple comparisons test). N.D. indicates that the data was not detected. d) The blood trickled from tail-tip in saline solution during 20 min bleeding test. See **Supplementary Video 4** for bleeding. e) Total blood loss during 20 min bleeding evaluated by hemoglobin concentration. Data represent the mean ± SEM (n=5, wild type; n=4-6, transplanted; n=4-5, sham; *, p<0.05; ***, p<0.001; Welch’s t-test). f) Shortening of time until bleeding stopped relative to maximum (20 min). Data represent the mean ± SEM (n=5, wild type; n=4-6, transplanted; n=4-5, sham; ***, p<0.001; Welch’s t-test). The activity of FVIII secreted from transplanted HLBO 2 and 5 months after transplantation, which evaluated by APTT shortening compared with normal human plasma, PBS, and rhFVIII. Data represent the mean ± SEM (n=3-5; **, p<0.01; ****, p<0.0001; one-way ANOVA with Dunnett’s multiple comparisons test compared to PBS injected mice).

## Discussion

Emerging evidence highlights the vital role of the endothelium with transcriptomic and anatomical features in exerting tissue-specific functions^67,68^, yet the self-organizing mechanisms governing their *in vitro* formation are poorly understood^69,70^. Despite recent advancements in organoid vascularization in the liver^24^, kidney^3,15^, and brain^71^, these studies utilized fully committed arterial endothelial cells for engineering endothelial networks in the organoids and therefore it remains unclear as to the endothelial cells’ specificity. Previous studies suggested that liver sinusoidal vessels originated from dual- or multi- progenitors including endocardium and vitelline vein connecting yolk sac to body of embryo^30,32,72^. The integrated single-cell transcriptome of endothelium from the heart, yolk sac, fetal liver and adult liver demonstrated that the expression of a group of angiogenic factors, such as VEGF- A and the OSM signaling pathway, is enhanced along the pseudo-time course from early-stage (4 PCW) yolk sac EC and endocardium with a marker expression profile typical to hepatic sinusoidal endothelium. In particular, OSM is a fetal liver development-associated signal released from hematopoietic cells that have migrated from the yolk sac to the fetal liver, suggesting that this regulated process is specific to liver angiogenesis^63^. This insight was enriched to derive LSECs from progenitors by modulating the OSM signaling in both 2D and 3D organoid approaches. Collectively, we have established a protocol to induce iLSEPs with high differentiation potential into LSECs in an OSM stimulation-dependent manner.

Despite recent progress in complex organoid culture, sustaining multicellular composition in culture remains a significant challenge. Here we developed the IMALI culture that allows the self-organization of sinusoid-like branched endothelial networks sprouted from iLSEP in liver organoids by introducing an air-liquid interface and leveraging multilayered patterns in culture^30,31,73^. The multilayered gel can support compartmentalization and compactization of tubular structure in testicular culture^46^. In our IMALI culture-based vascularization, two different types of ECs in a separated gel layer undergo an aggregation process to achieve organoid formation with hierarchical vessels. While CD31^+^CD32b^-^ ECs form a primitive plexus-like structure, CD32b^+^FVIII^+^ ECs migrates and sprouts towards the more oxygenated upper layer. iLSEPs differentiated not only into sinusoidal ECs but also into THBD^+^ central vein-like endothelial cells, implying that the incorporating multiple endothelial progenitor cells is propulsive in constructing endothelial hierarchy likely through combination of vasculogenesis and angiogenesis processes^74^.

Angiocrine signals such as Wnt2 and RSPO3 released from LSECs contribute to hepatocyte maturation^12,16,63,64^. Further understanding of the spatiotemporal interactions between cells is a crucial roadmap to elucidate the processes that achieve the architecture and function of the liver^75^. However, there is a lack of *in vitro* models that reflects endothelial diversity to track the developmental process. Our human organoid model system delineated key insights for organ-specialized vessels that involves complex multicellular interactions. For example, we noted the emergence of both GLS2^+^ and GS^+^ hepatocyte subsets in the IMALI- based liver organoid model. Indeed, the formation of CD32b^+^FVIII^+^LYVE1^+^ liver sinusoid-like endothelium enhanced peri-central hepatocytic metabolic enzyme expression and activity. Endothelial cell-specific knockdown strategy in HLBO indicated Wnt2 mediated crosstalk is essential for gaining hepatocyte differentiation and endothelial branching. Future exploration of the other reciprocal angiocrine crosstalk among endothelial and non-endothelial subsets in HLBO system will inform the divergent roles of hepatic vasculature in supporting liver development in humans.

Our sinusoid-bearing HLBO produces a variety of coagulation factors, especially FVIII, which can persist for about 40 days. Given that there are no available human protein sources for some classes of coagulation factors, such as FV and FXI, functional coagulation factors secreted from human organoids might be an authentic source for producing critical proteins for the development of diagnostics and therapeutics for disorders affecting the coagulation cascade. This culture supernatant harbors stable activity even in the presence of inhibitors, typically observed in 20-30% of severe hemophilia A patients^76^, and the multi- production of coagulation factors such as FII, FVII, and FX may offer an advantage for the treatment of patients at high risk of developing inhibitors to coagulation factors.

*In vivo*, transplanted HLBO was connected to the host blood vessels, reconstructed human blood vessels fulfilling several key sinusoidal characteristics including acetylated low-density lipoprotein endocytic capability, permeability and supplied active FVIII up to 20 weeks in the FVIII KO mice. Recently, clinical trials of gene therapy using adeno-associated vectors have been underway, but the immunogenicity and persistence of viral FVIII gene transfer remain unclear^77^. In recent PSC-derived LSEC transplantation efforts, pre-conditioning with monocrotaline is necessary for engraftment^26,28,72,78^. In another *in vivo* transplantation study, Son et al. reported that attempted to improve coagulopathy for up to 2 weeks^79^. Our *in vivo* transplantation successfully corrected the coagulation disorder phenotype for up to 8 weeks without requiring toxin exposure. Thus, cellular- or organoid-based regenerative technology offers an alternate way of developing new therapeutic concepts for coagulation disorders.

## SUPPLEMENTARY FIGURE LEGENDS

**Extended Data Fig. 1:**
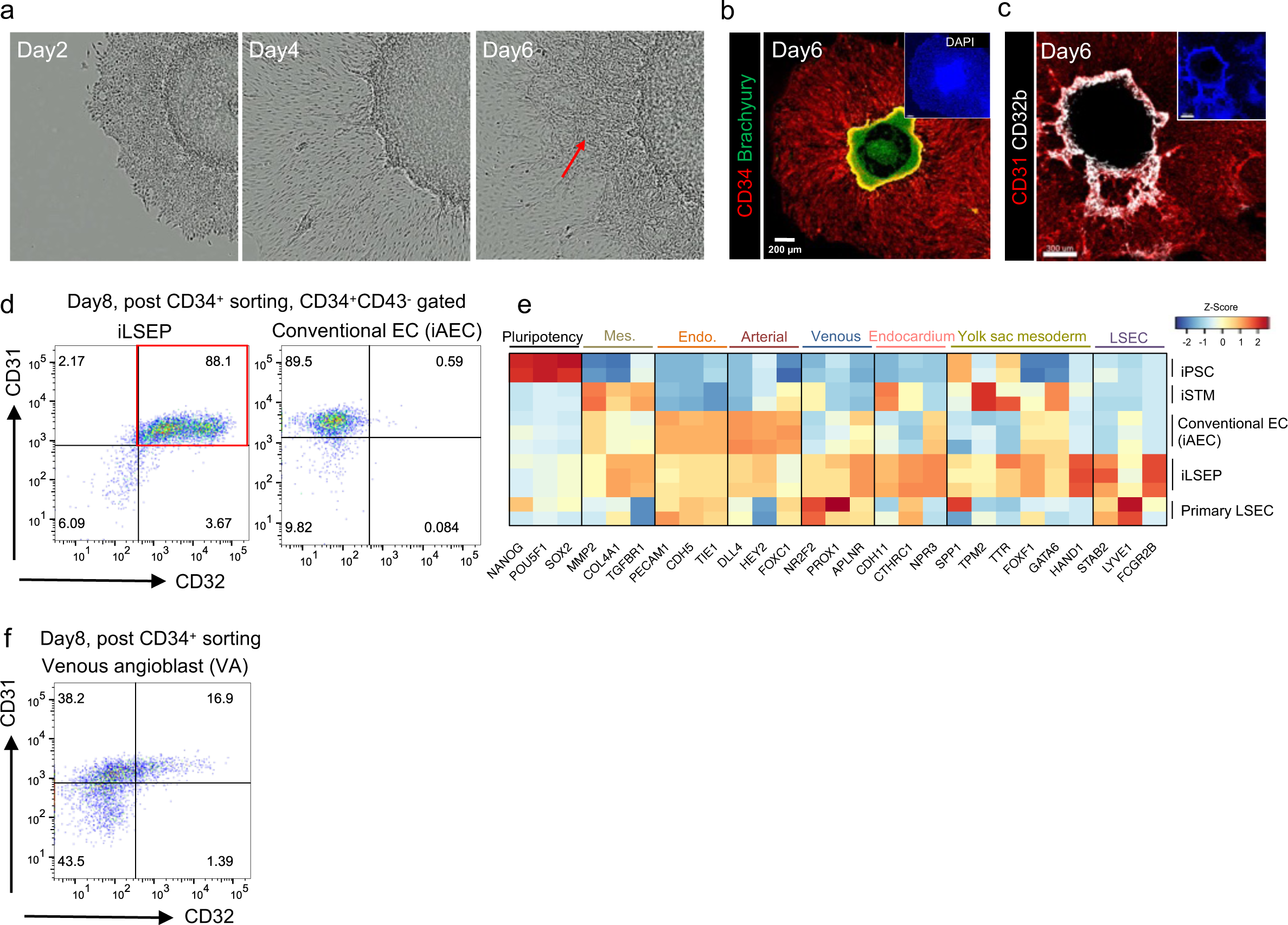
Directed differentiation and characterization of iLSEP. a) Time-lapse morphological representation of differentiation of iLSEP in micropatterned colony. Red arrows indicate expanding CD34^+^ endothelial regions (together with **Extended Data Fig. 1b**). b) Immunofluorescent staining for CD34 and Brachyury at day6. c) Immunofluorescent staining for CD31 and CD32b at day6. d) Scatter dot-plot of the flow cytometry analysis of CD34/CD32 expression in iLSEP and iAEC after CD34^+^ purification at day 8. e) Heatmap of iPSC, iSTM, conventional EC (iAEC), iLSEP, and primary LSEC showing lineage marker gene set. Mes.: Mesenchyme; Endo.: pan-endothelium; LSEC: liver sinusoidal endothelial cell. f) Scatter dot-plot of the flow cytometry analysis of CD31/CD32 expression in CD34^+^CD43^-^ VA at day8.

**Extended Data Fig. 2:**
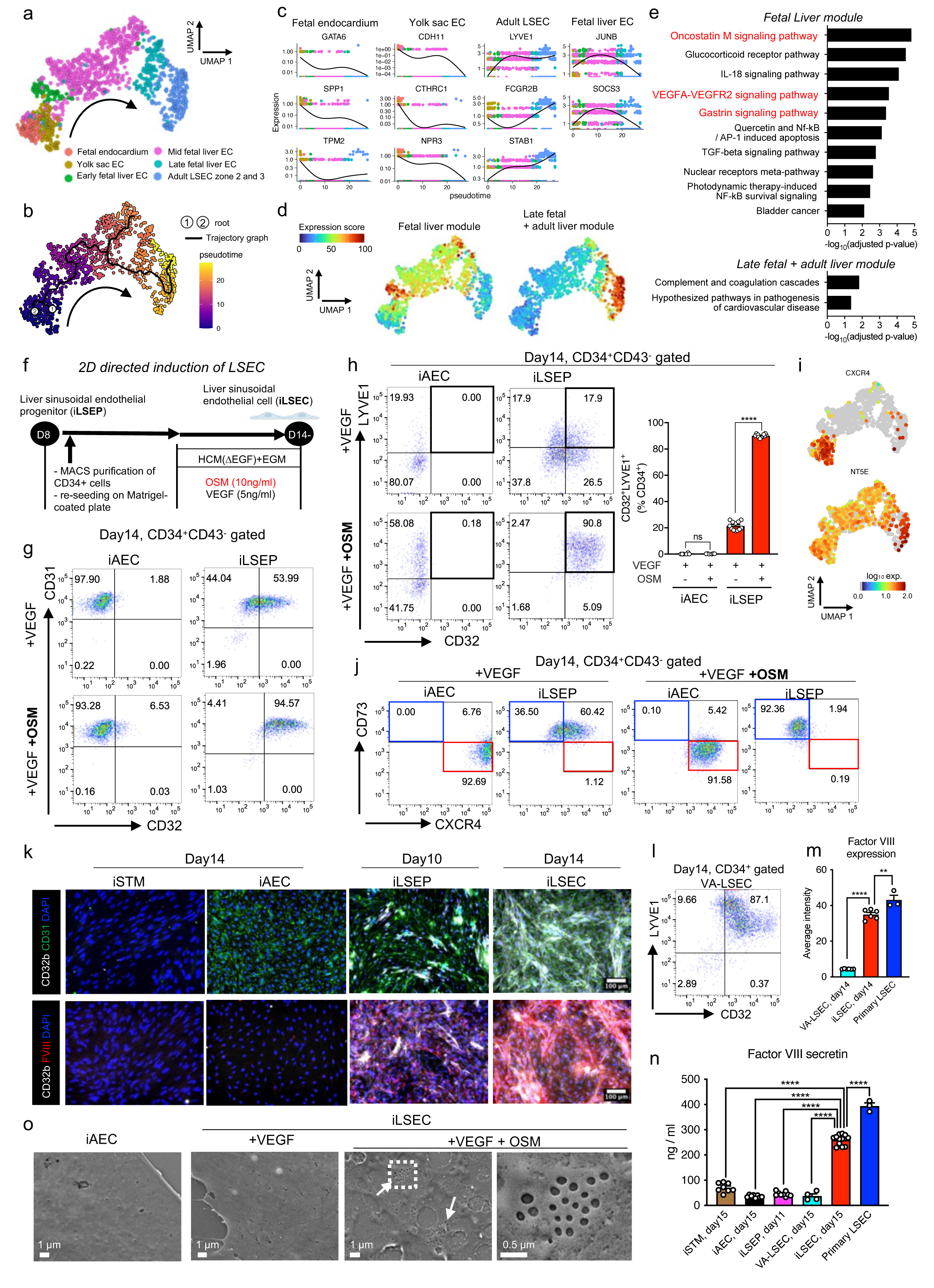
Induction of LSEPs into LSECs by OSM, guided by single-cell analysis of endothelial development. a) Integrated uniform manifold approximation and projection (UMAP) plot of ECs in fetal and adult organs showing yolk sac EC (4 post conception weeks (PCW)), endocardium in fetal heart (4 PCW), fetal liver EC (Early; 7 PCW, Mid; 8-12 PCW and Late; 13-17 PCW), and LSEC zone 2 and 3 in adult liver. The black arrow curve represents the direction of the trajectory based on pseudo-temporal analysis (together with **Extended Data Fig. 2b**). b) The pseudo-time trajectory of endothelial development to LSEC in adult liver. c) Pseudo-kinetic plots of the expression of markers in different organ groups. d) The aggregate expression level of all genes in fetal (early, mid, and late) liver module and late fetal + adult liver module projected on UMAP. e) Top significantly enriched pathways on dynamics of fetal liver module and late fetal + adult liver module. f) The schema of iPSC-derived LSEC differentiation via LSEP. g) Scatter dot-plot of the flow cytometry analysis of CD31/CD32 expression in CD34^+^CD43^-^ iAEC and iLSEP at day14 under +VEGF and +VEGF+OSM conditions. h) Scatter dot-plots of the flow cytometry analysis of LYVE1/CD32 expression (left). Quantification of CD32^+^LYVE1^+^ cell fractions in CD34^+^CD43^-^ cells (right). Data represent the mean ± SEM (n=6, iAEC; n=12; iLSEP; ****, p<0.0001; ns, not significant; one-way ANOVA with Tukey’s multiple comparisons test). i) The feature plots showing the expression of arterial (CXCR4) and venous (NT5E coding CD73 protein) EC markers in integrated single-cell gene expression data. Gray color indicates no expression. j) Scatter dot-plots of the flow cytometry analysis of CD73/CXCR4 expression. k) Immunofluorescent staining for CD32b/CD31 (upper) and CD32b/FVIII (lower) at day 12 (iSTM, iAEC and iLSEP) and day 16 (iLSEC). l) Scatter dot-plot of the flow cytometry analysis of LYVE1/CD32 expression in VA-derived LSEC (VA-LSEC) at day14. m) Comparison of FVIII protein expression quantified from images of immunofluorescent staining for FVIII/CD32b. n) Comparison of secreted FVIII concentration in culture supernatants among iPS-derived STM, endothelial cells, and primary LSEC. Data represent the mean ± SEM (n=3-12; ****, p<0.0001; one-way ANOVA with Dunnett’s multiple comparisons test compared to iLSEC, day15). o) Scanning electron microscopy (SEM) of iLSEC (+VEGF and +VEGF, +OSM) and iAEC. Arrowheads indicate the clusters of fenestrae on the cell surface. The right panel is an enlarged image of the dashed square region of iLSEC (+VEGF, +OSM).

**Extended Data Fig. 3:**
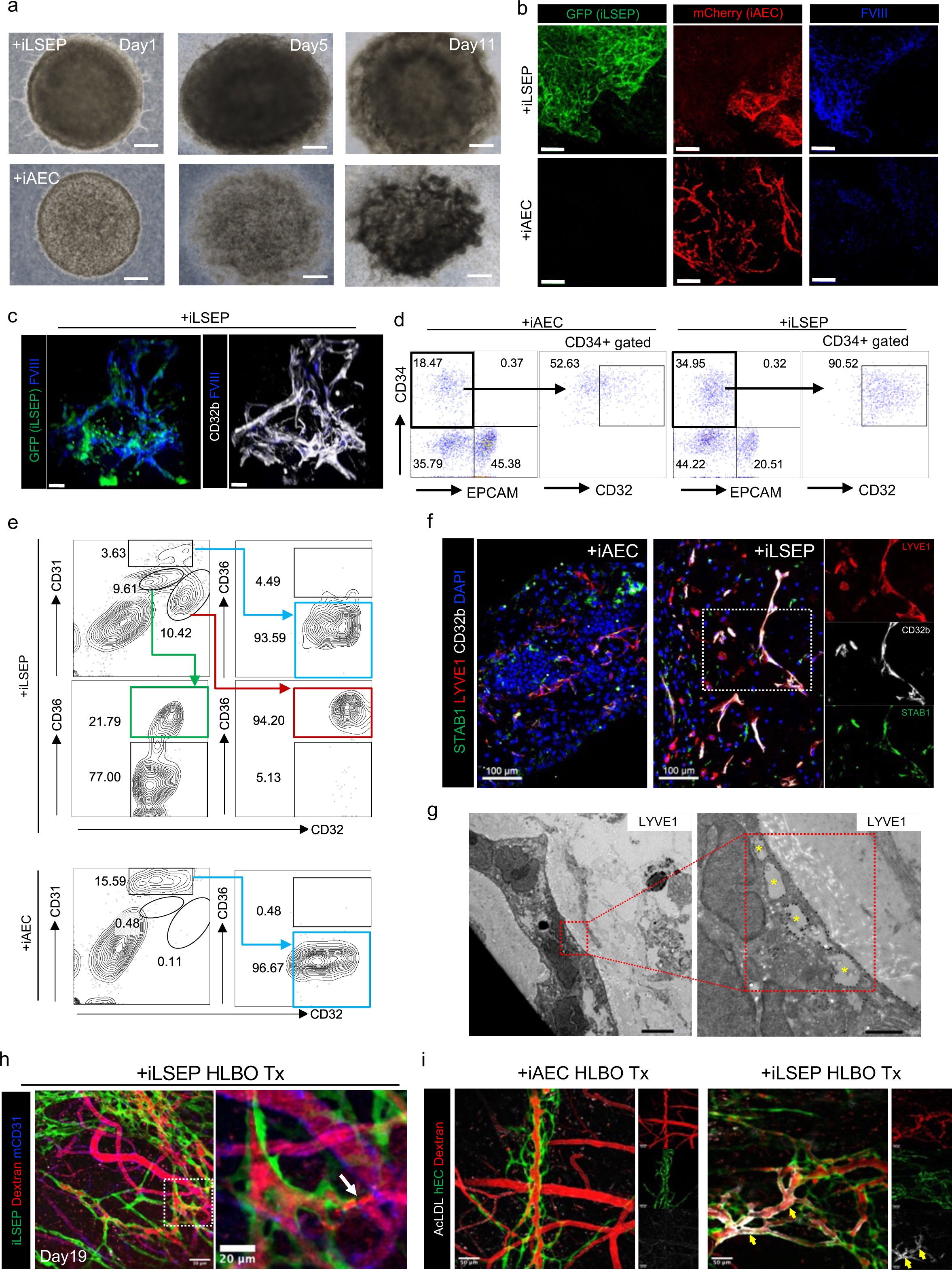
Characterization of endothelial lineages in self-organizing liver bud organoids. a) Bright-field image of organoids at different time points after mixing cells (day 1, 5, and 11). Scale bar indicates 500 µm. b) Whole-mount single fluorescent channel images of HLBOs including GFP^+^ iLSEP and mCherry^+^ iAEC with immunofluorescent staining for FVIII at day 11. Scale bar indicates 200 µm. c) High-magnification whole mount image of HLBO including GFP^+^ iLSEP with immunofluorescent staining for CD32b and FVIII at day 11. Scale bar indicates 30 µm. d) Scatter dot-plots of the flow cytometry analysis of CD34^+^ endothelial cells and EPCAM^+^ (CD326^+^) epithelial cells in HLBOs at day11. The right panels display CD32 expression in endothelial cells gated by CD34. e) Scatter dot-plots of the flow cytometry analysis of CD31, CD32, and CD36 expression of HLBOs at day11. The panel of CD32 and CD36 was gated by CD31^high^ for both HLBOs, and CD32^high^CD31^low^ and CD32^low^CD31^low^ only for +iLSEP HLBO. f) Cross-sectioning image of HLBOs with immunofluorescent staining for LYVE1/CD32b/STAB1 at day 11. Right panels are enlarged single fluorescent channel images of the dashed square region. g) Transmission immunoelectron microscopy of ultra-thin frozen sections of +iLSEP HLBO labeled with anti-LYVE1 antibody. Right panel is enlarged image of the red dashed square region of left panel. Yellow asterisks indicate pore structures between LYVE1^+^ endothelial cells. Scale bar indicates 5 µm (left) and 1 µm (right). h) Intravital fluorescence microscopy imaging of the +iLSEP HLBO transplanted mice. The organoid-derived human vessels were observed by GFP^+^ iLSEP. Host mice vessels and blood flow are visualized using a mice-specific anti-CD31 antibody (mCD31) and 2,000 kDa tetramethylrhodamine-dextran, respectively. Right panels are enlarged image of the dashed square region. White arrow marks the point of connection between the human and mouse blood vessels. i) Alexa Fluor 647 (AF647)-conjugated AcLDL uptake in human blood vessels with blood flow (2,000 kDa tetramethylrhodamine-dextran). Right panels are single fluorescent channel images. Yellow arrowheads indicate regions of AF647-AcLDL merged human blood vessels. Scale bar indicates 50 µm.

**Extended Data Fig. 4:**
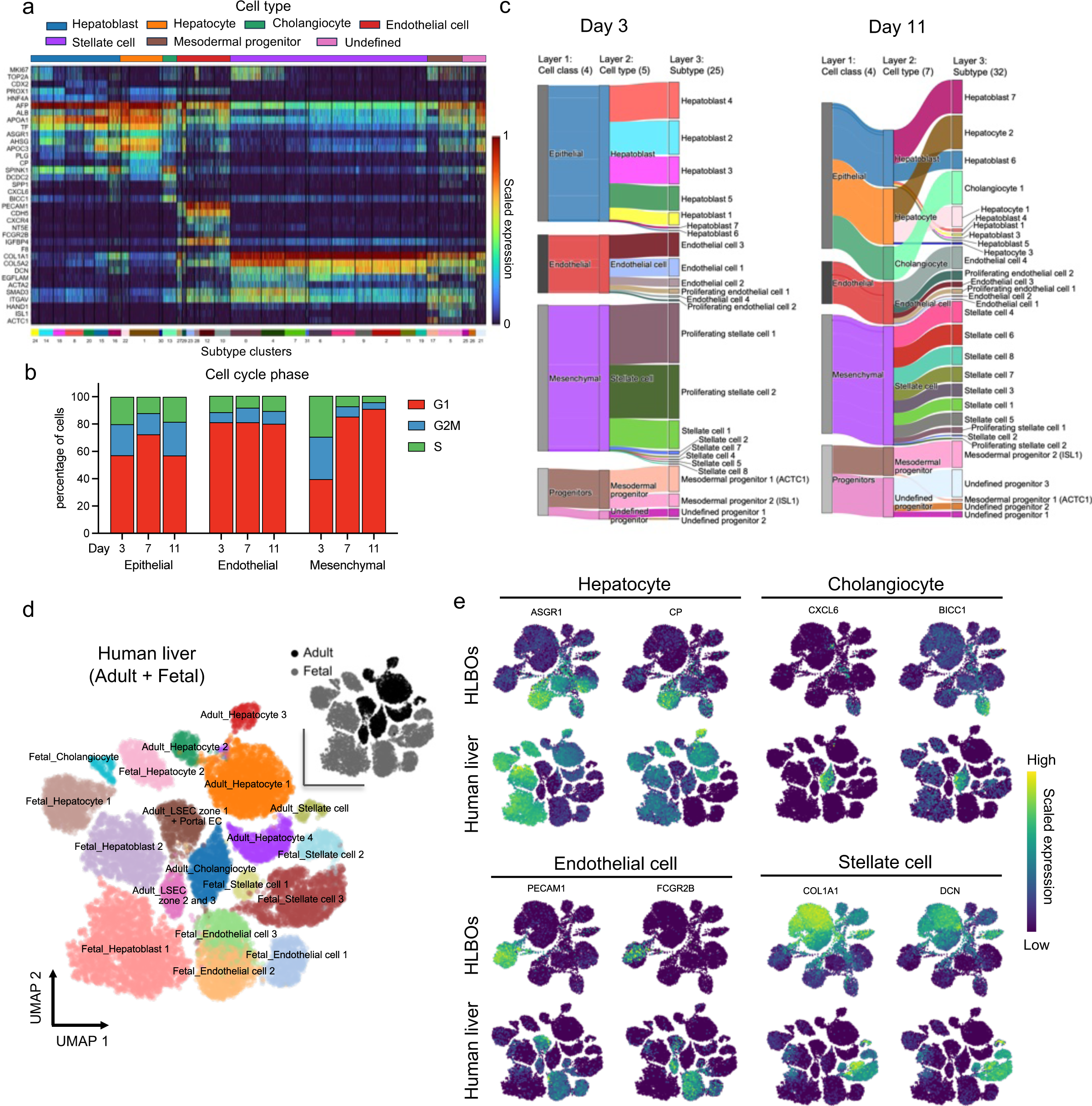
scRNA-seq profiling of HLBOs and primary human liver. a) Heatmap showing marker gene expression for cell subtype layer group in the HLBO+iLSEP dataset. b) Proportion of cell cycle phase per cell class in each time series. c) Sankey diagram for hierarchical overview of cell composition in HLBO+iLSEP. The hierarchical structure consists of three annotation levels: cell class (Layer 1), which is the upper domain grouping cell types, cell type (Layer 2), and subtype (Layer 3), and the numbers in parentheses indicate the total number of annotations in each layer. d) UMAP of integrated primary liver dataset including fetal and adult stage colored by cell type. e) UMAPs showing common cell type marker expression between primary liver and HLBO+iLSEP.

**Extended Data Fig. 5:**
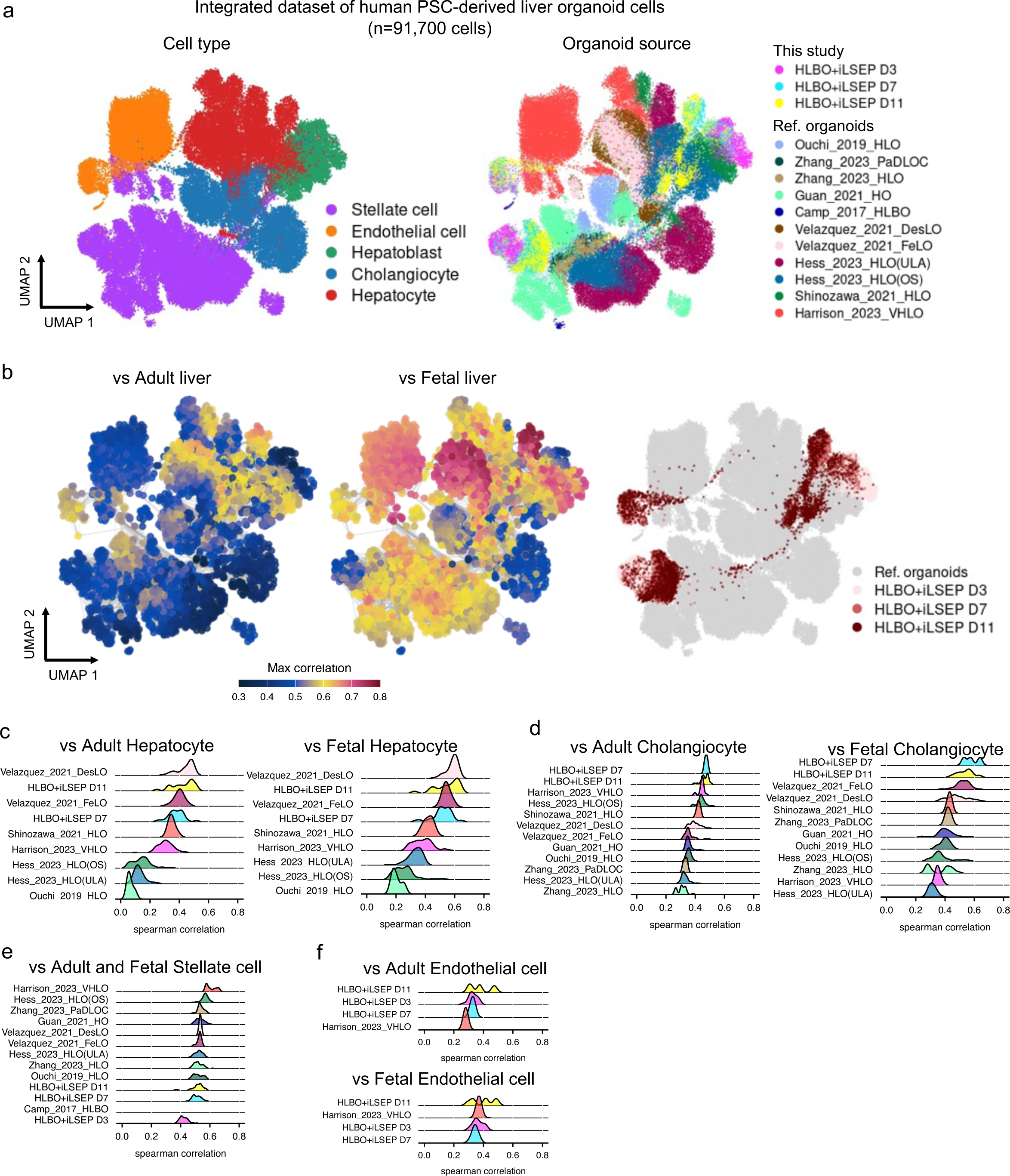
Comparative analysis of HLBO with published human organoid datasets. a) UMAPs for human PSC-derived liver organoid cell atlas colored by cell type and organoid source. b) UMAPs display the maximum spearman correlation of (left) adult liver and (right) fetal liver dataset. The right UMAP highlights HLBO+iLSEP cells. c-f) Ridge plots displaying spearman correlations for each organoid dataset against c) adult or fetal hepatocytes, d) adult or fetal cholangiocytes, e) both adult and fetal stellate cells, and f) adult or fetal endothelial cells. Organoid groups that did not contain enough cells to show data distribution are not plotted.

**Extended Data Fig. 6:**
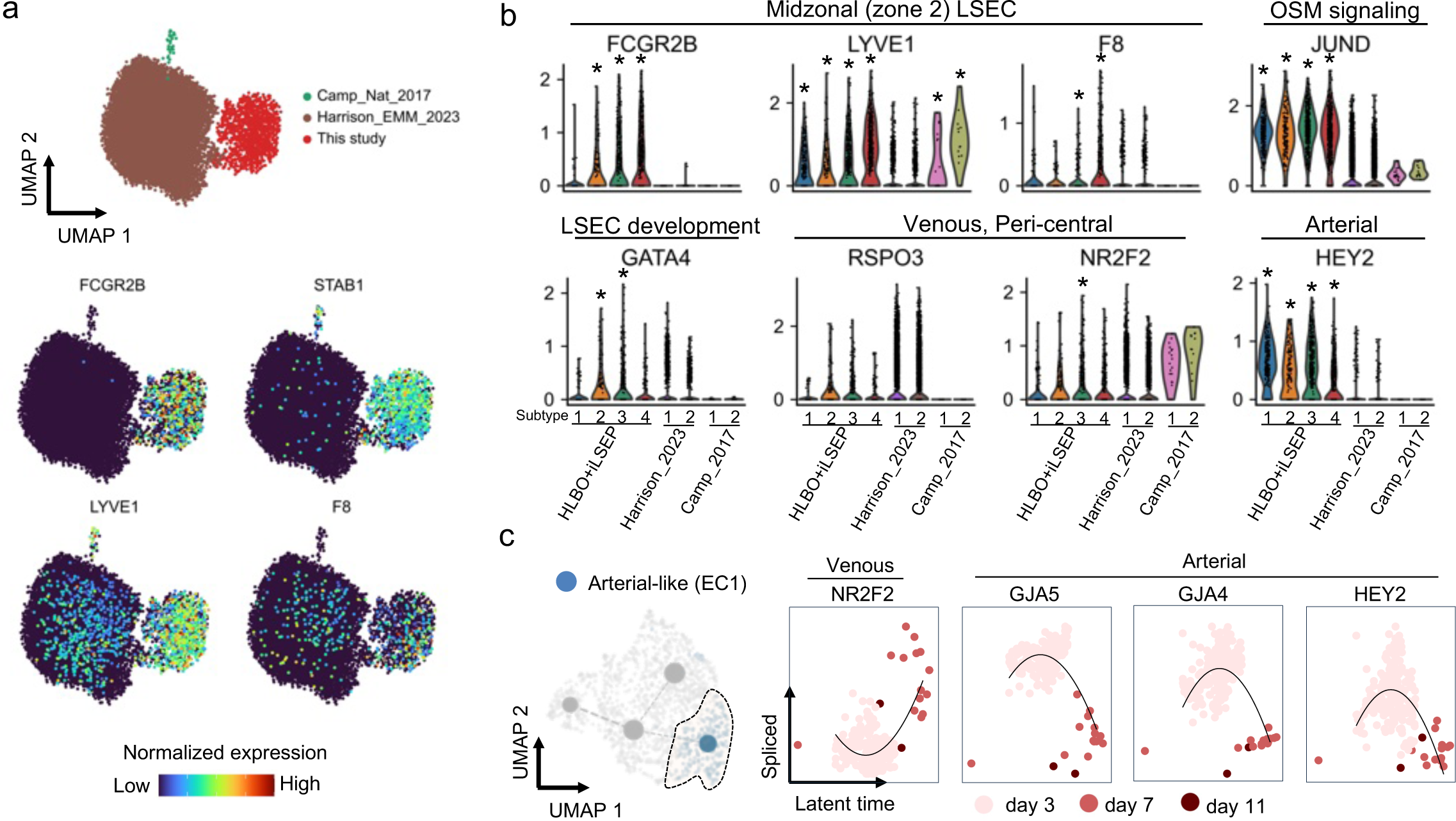
Single-cell characterization of endothelial cell in liver organoids a) UMAPs showing LSEC (zone 2 and 3) marker gene expression (together with **Extended Data Fig. 2c**), whereas organoids are color-coded by data resource. b) Violin plots showing differentiation signature expression by endothelial subset. c) UMAP showing extracted subpopulation of arterial-like endothelial cell (EC1) (left) and Scatter plots showing latent time versus spliced gene expression of arterial markers (right). Black line on scatter plot is linear regression fit line to data points to represent trends of change.

**Extended Data Fig. 7:**
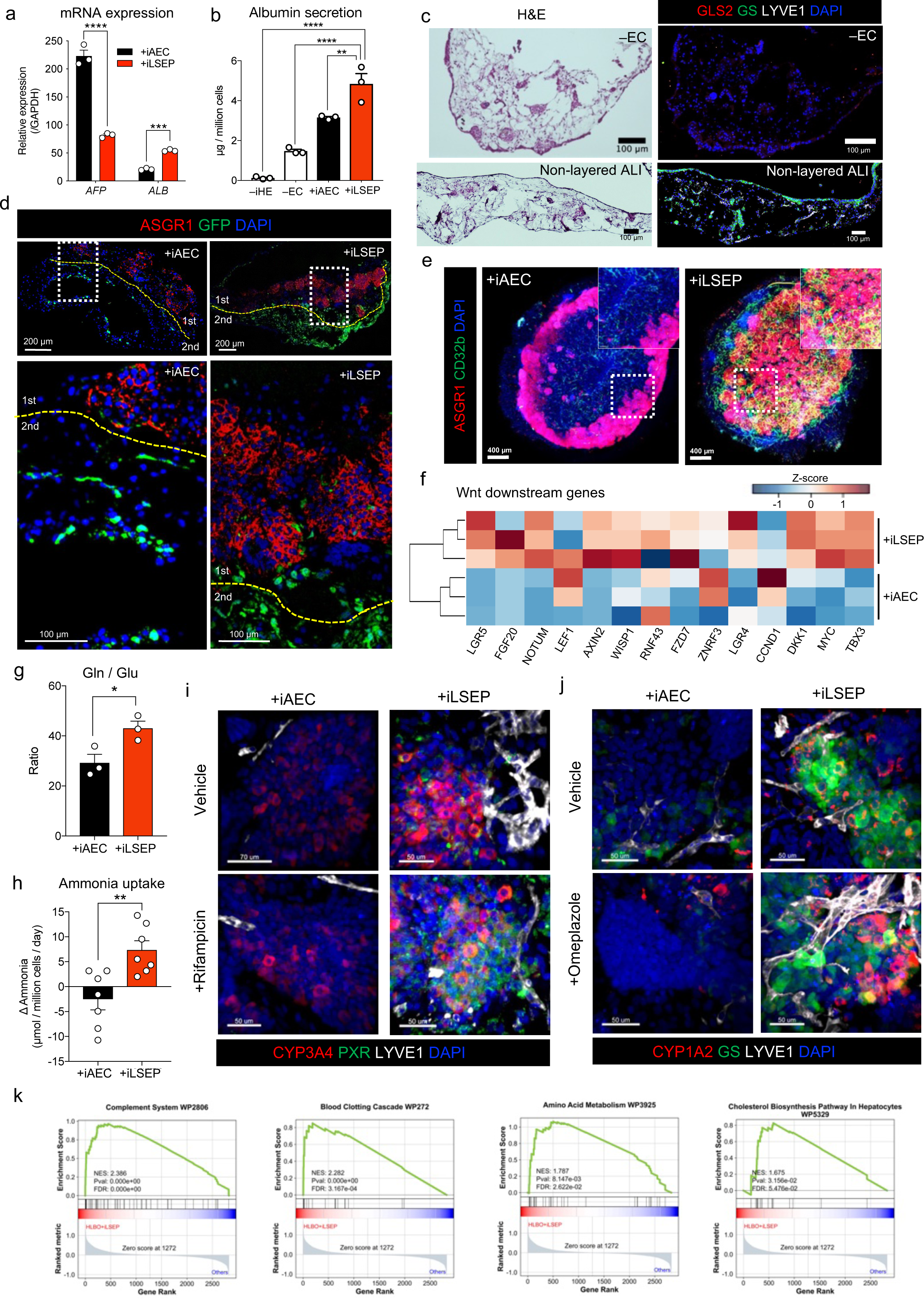
Analysis of hepatic functionalization and architecture in HLBOs. a) qRT-PCR analysis of AFP and ALB. Data represent the mean ± SEM (n=3; ***, p<0.001; ****, p<0.0001; Welch’s t-test). b) Measurement of albumin in organoid supernatants at day 11. Data represent the mean ± SEM (n=3; **, p<0.01; ****, p<0.0001; one-way ANOVA with Dunnett’s multiple comparisons test compared to +iLSEP HLBO). c) Cross-sectioning image of HLBO without EC (-EC) with H&E staining (left). Distribution of hepatocytes with different zonal marker protein identified by immunofluorescent staining for GS (peri-central) and GLS2 (pan-lobular) in HLBOs (right). d) Cross-sectioning images showing distribution of hepatocytes identified by immunofluorescent staining for ASGR1 and GFP^+^ iAECs or iLSEPs embedded in a second cell layer. Lower panels are enlarged images of the white dashed square region. The yellow dashed lines indicate the boundaries of the gel layer in organoid generation as illustrated in Fig. 1a. e) Localization of ASGR1^+^ hepatocyte and CD32b^+^ LSEC determined by whole-mount immunofluorescent staining of HLBOs. f) The heatmap showing expression of the Wnt downstream target gene set. g) Measurement of glutamine / glutamate ratio in HLBO supernatants at day11. Data represent the mean ± SEM (n=3; *, p<0.05; Welch’s t-test). h) Determination of ammonia uptake from the medium in HLBOs at day 11. Ammonia change in 24 hours was quantified by the difference between the HLBO supernatant and the culture medium. Data represent the mean ± SEM (n=7; **, p<0.01; Welch’s t-test). i-j) Whole-mount immunostaining images of HLBO with/without CYP inducers stained for CYP3A4/PXR/LYVE1 and CYP1A2/GS/LYVE1. Right panels are fluorescence channel split images of LYVE1 combined with each hepatocyte markers in +iLSEP HLBO. j) Representative enrichment plots compared between hepatocyte in +iLSEP HLBO and those in other liver organoids (9 publicly available organoids) of integrated dataset.

**Extended Data Fig. 8:**
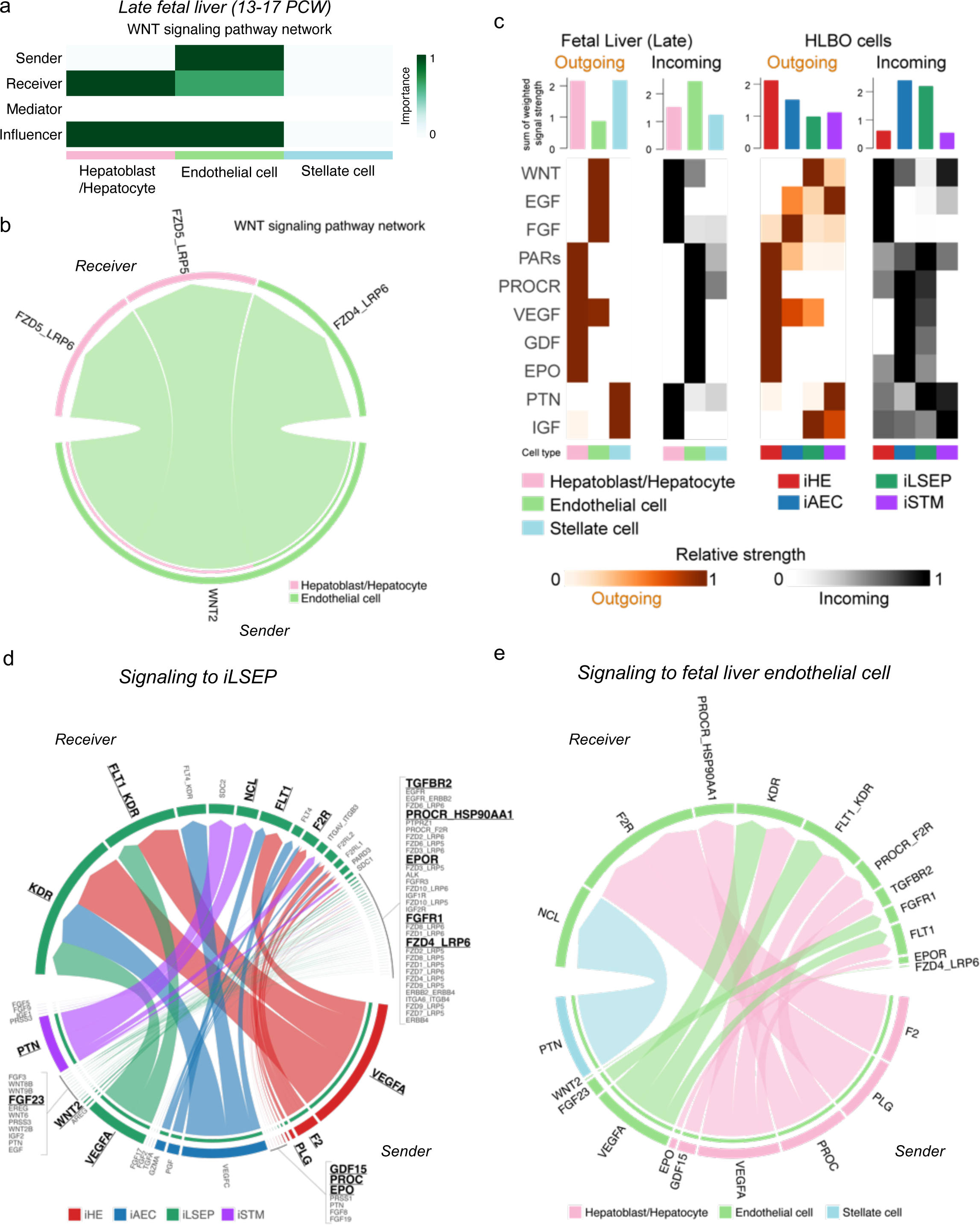
Inter-cellular lineage interaction by ligand-receptor analysis. a) The heatmap visualizing the relative importance of each cell type of late fetal liver (13- 17 PCW) based on the centrality score of WNT signaling pathway. b) The chord diagram describing significant ligand-receptor pairs involving WNT signaling within fetal liver (p<0.05). c) The heatmap and bar plot summarizing outgoing or incoming signaling strength across cell types. Color scale of the heatmap represents the relative signaling strength of each pathway. Top bar plot shows total strength of all pathways displayed in the heatmap for each cell type. d) The chord diagram describing significant ligand-receptor pairs involved all signals sending to iLSEP (p<0.05). The ligand and receptor names that overlap with intercellular communication in fetal liver are highlighted. e) The chord diagram describing significant ligand-receptor pairs involved all signals sending to fetal liver endothelial cell (p<0.05).

**Extended Data Fig. 9:**
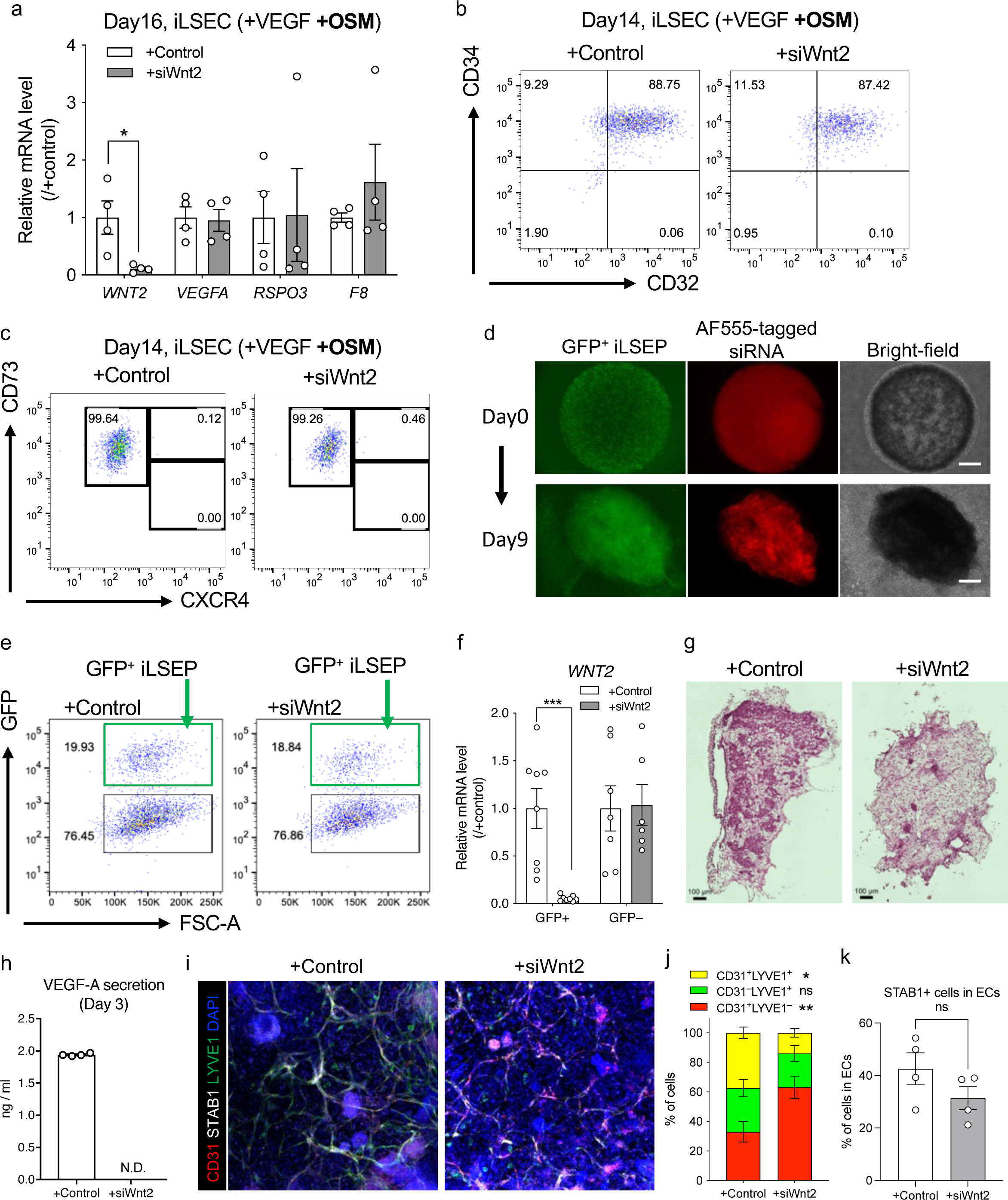
Evaluation of iLSEP-specific siRNA knockdown. a) qRT-PCR analysis of *WNT2*, *VEGFA*, *RSPO3*, and *F8*. Data represent the mean ± SEM (n=4; *, p<0.05; multiple t-test with controlling FDR). b-c) Scatter dot-plot of the flow cytometry analysis of CD34/CD32 and CD73/CXCR4 expression in CD34^+^CD43^-^ iLSEC with siRNA treatment at day14. d) Observation of fluorescent reporters with bright-field image to track GFP^+^ iLSEP and AlexaFlour555 (AF555)-tagged siRNA embedded in the same layer of multilayered gel. Scale bar indicates 500 µm. For at least 9 days, AF555-tagged siRNA remained in the packed gel or inside the organoids. e) Scatter dot-plot of the flow cytometry analysis of whole HLBO cells for separation of GFP-tagged iLSEP in second layer and GFP-negative cells in first layer in HLBO by FACSAria Fusion cell sorter. f) qRT-PCR analysis of Wnt2 to validate GFP-tagged iLSEP specific knockdown efficiency. WNT2 mRNA expression levels were determined relative to the negative control siRNA treated for each GFP^+^ and GFP^-^ fraction isolated. Data represent the mean ± SEM (n=6- 8; ***, p<0.001; multiple t-test with Holm-Sidak correction). g) Cross-sectional H&E staining image of HLBOs with siRNA treatment. h) Measurement of VEGF-A in siRNA-treated HLBO supernatants at day3. Data represent the mean ± SEM (n=4; ND, Not detected). i) Whole-mount image of siRNA-treated HLBOs with immunofluorescent staining for CD31, STAB1 and LYVE1. j) Quantification of the percentage of cells expressing CD31 or LYVE1 alone or co-expressing CD31 and LYVE1. Data represent the mean ± SEM (n=4; *, p<0.05; **, p<0.01; two-way ANOVA with Sidak’s multiple comparison test). k) Quantification of the percentage of STAB1 co-espressing endothelial cells (CD31^+^ or LYVE1^+^). Data represent the mean ± SEM (n=4; ns, not significant; Welch’s t-test)

**Extended Data Fig. 10:**
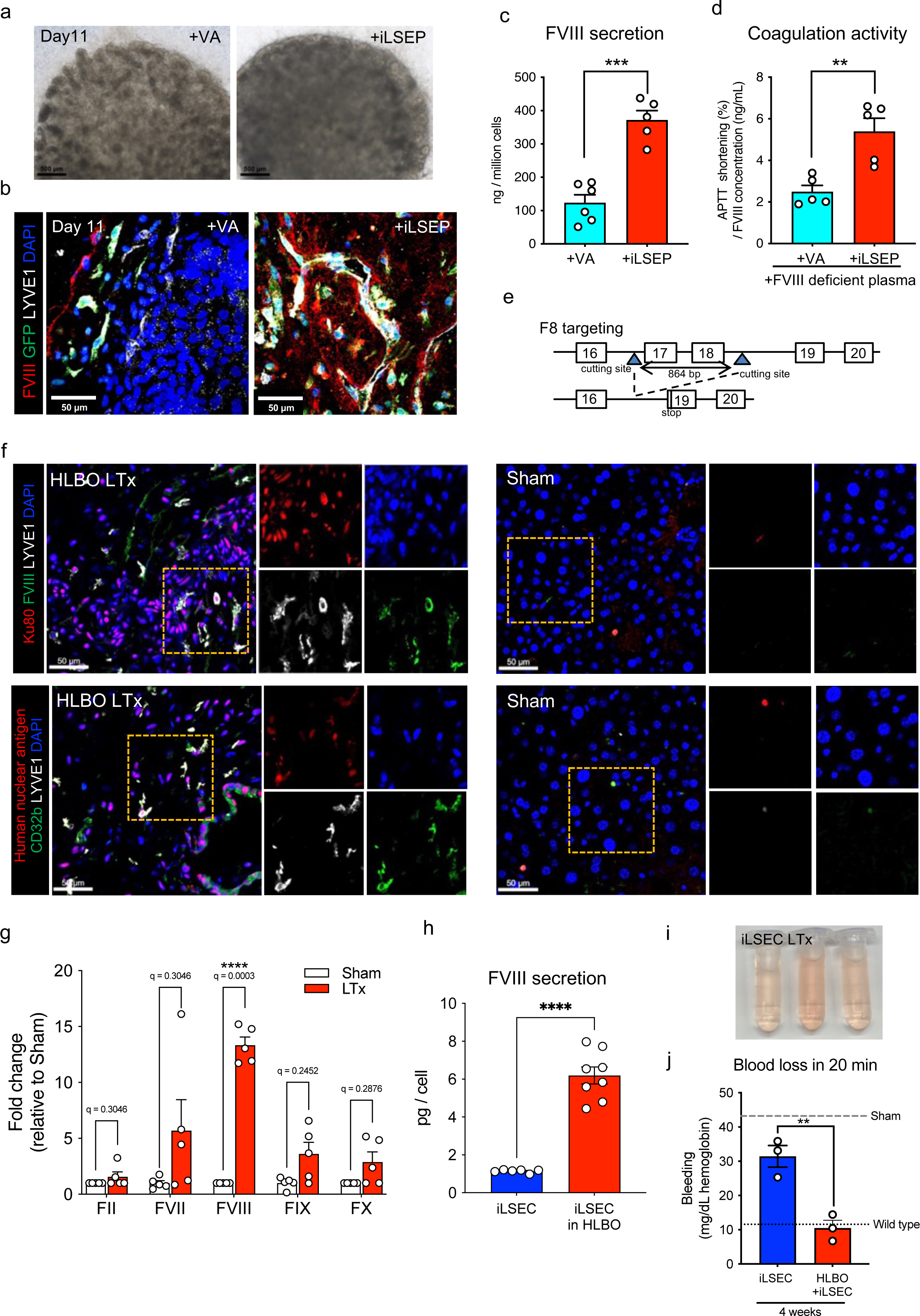
Assessment of Factor VIII in human fetal livers and HLBOs incorporating VA or transplanted into FVIII knockout mice. a) Bright-field image of organoids at day11. Scale bar indicates 500 µm. b) Cross-sectioning image of HLBOs with immunofluorescent staining for FVIII and GFP- tagged VA or iLSEP embedded in a second cell layer, and LYVE1 at day 11. c) FVIII concentration in organoid supernatants at day 11. Data represent the mean ± SEM (n=5-6; ***, p<0.001; Welch’s t-test). d) APTT shortening activity of organoid supernatants at day 11 with FVIII deficient plasma normalized by FVIII concentration. Data represent the mean ± SEM (n=5; **, p<0.01; Welch’s t-test) e) The schema of FVIII gene editing to establish the immunodeficient NS-IL2rg KO/ FVIII KO hemi (NSI-FVIII KO) mice. f) Cross-sectioning image of HLBO graft and mice liver tissue with immunofluorescent staining for human FVIII, LYVE1, CD32b, and human cell marker (Ku80 or human nuclear antigen). Right figure for each group is enlarged single channel images of the orange dashed square region in left figure. g) Measurement of HLBO derived human coagulation factors circulating in mouse blood 8 weeks after liver transplantation (LTx) (n=5, both Sham and transplanted; **, one-way ANOVA with Tukey’s multiple comparisons test for statistical evaluation). h) FVIII concentration in culture supernatants on day 7 after re-seeding (iLSEC) or self-organization (iLSEC in HLBO). Values were normalized by the number of LSEC. Data represent the mean ± SEM (n=6-8; ****, p<0.0001; Welch’s t-test). i) The blood trickled from tail-tip in saline solution during 20 min bleeding test. j) Total blood loss during 20 min bleeding evaluated by hemoglobin concentration. Data represent the mean ± SEM (n=3; **, p<0.01; Welch’s t-test).

**Supplementary Video 1: Whole-mount 3D reconstructed movie of +iLSEP HLBO with immunofluorescent staining for CD32b (green) and ASGR1 (red) with DAPI (blue) at day11, related to** Fig. 1c.

**Supplementary Video 2: Time-lapse movie of 70 kDa or 2,000 kDa tetramethylrhodamine-dextran visualized-blood flow in GFP^+^ organoid-derived human vessels for 60 min, related to** Fig. 1i.

**Supplementary Video 3: *in vivo* hemophilia A correction by the HLBO supernatant injection, related to** Fig. 5i, j.

**Supplementary Video 4: *in vivo* hemophilia A correction 8 weeks after orthotopic transplantation of HLBO, related to** Fig. 6d**-f**.

**Supplementary Table 1: Significant marker statistics for each group in integrated single-cell transcriptome of endothelial cells in adult liver, fetal liver, yolk sac, and fetal heart (q value<0.05), related to Extended Data** Fig. 2a-c.

**Supplementary Table 2: Top 100 marker gene list of all endothelial groups in integrated single-cell transcriptome, related to Extended Data** Fig. 2a-c.

**Supplementary Table 3: The list of genes in modules that changed in trajectory to LSEC zone 2 and 3, related to Extended Data** Fig. 2d, e.

**Supplementary Table 4: The detailed list of significantly enriched pathways on dynamics of gene module along the trajectory to LSEC zone 2 and 3, related to Extended Data** Fig. 2e.

**Supplementary Table 5: Significant marker statistics for each group in integrated single-cell transcriptome of +iLSEP HLBO day 3, 7, and 11 (qvals_adj<0.05 and log_fold_change>2), related to** Fig. 2a.

**Supplementary Table 6: Description of collected data for construction of a dataset integrated with published human liver organoids, related to Extended Data** Fig. 5.

**Supplementary Table 7: Significant marker statistics for each endothelial subpopulation in HLBO (pvals_adj<0.05), related to** Fig. 2b.

## Methods

### Single-cell transcriptome data integration and analysis for human liver endothelial development

The public single-cell transcriptome data were obtained as follows. For fetal heart and adult liver, the Unique. Molecular Identifiers (UMI) count matrix and barcode information deposited in GSE106118 and GSE115469 of Gene Expression Omnibus (GEO) were used. For fetal liver and yolk sac, fastq files were obtained from E-MTAB-7407 of ArrayExpress, and the sequence reads were processed using the count pipeline implemented in Cell Ranger Software Suite version 2.0 (10x Genomics). Raw reads were aligned to the GRCh38 reference genome and filtered for creating UMI matrices with default parameters.

All UMI count matrices were subjected to cell and gene filtering and data scaling using the Seurat v3 package implemented in R. Cells which have unique feature counts less than 500 or mitochondrial counts more than 20% were removed. Genes which were expressed in fewer than 3 cells were also removed. The data was normalized using default global-scaling method. Subsequently, cell populations annotated as endothelial cells were extracted as input into downstream analysis.

Data integration, clustering, visualization, pseudo-time trajectory analysis and differentially expression analysis were performed in Monocle 3 Software (1.0.0). Scaled endothelial cell data from fetal heart, liver, yolk sac, and adult liver were integrated through batch correction using mutual nearest neighbor alignment with dimensionally reduction loaded 50 principal components for UMAP visualization. A principle graph was fitted on UMAP using the “learn_graph()” function to represent the path of the generation process. The root node of pseudo-time trajectory was set programmatically from the population of the earliest developmental stage (4PCW) using the “get_earliest_principal_node()” function.

Modules of co-regulated genes among the clusters was identified using “find_gene_modules()” function. Pathway enrichment analysis of the identified gene modules was performed using g:Profiler.

### hiPSC Culture

The human iPSC lines 1383D2, 625A4 (GFP reporter line) and 511-3E (mCherry reporter line) were kindly provided by the Center for iPS Cell Research and Application, Kyoto University. All iPSCs used herein had normal karyotypes (data not shown). All iPSCs were cultured on laminin 511 E8-fragment-coated (iMatrix-511; Nippi, Tokyo, Japan) dishes in StemFit AK02N (Ajinomoto, Tokyo, Japan).

### Inducded liver sinusoidal endothelial progenitor (iLSEP) induction

For differentiation towards iLSEP, the differentiation protocol as previously described^36,37^ was modified as follows. Briefly, undifferentiated PSC colonies were first disseminated onto laminin 511 E8- fragment-coated 10cm dishes in StemFit AK02N at a density of 30 clusters. When an individual colony grew to approximately 700-800 μm in diameter, differentiation was started by replacing with Essential 8 medium (Gibco) containing BMP4 (80 ng/mL), VEGF-A (80 ng/mL) and with or without CHIR99021 (2 µM) or IWP2 (2 µM) or LDN193189 (2 µM) for 2 days. On day 2, micropatterned colony including PS and YSM was formed, and the medium was replaced with Essential 6 medium (Gibco) containing VEGF-A (80 ng/mL), SCF (50 ng/ml), bFGF (25 ng/ml), and the TGFb inhibitor SB431542 (2 µM) for hemangioblast induction. On day 4, the medium was replaced with StemPro-34 serum-free medium (Gibco) containing 2 mM glutamax (Invitrogen), VEGF-A (50 ng/ml), SCF (50 ng/ml), Flt-3 ligand (10 ng/ml), IL-3 (10 ng/ml), and TPO (5 ng/ml) for transition to hemogenic endothelial lineage.

On day 6, CD34^+^ endothelial cells were formed at the outer edge of colony and the medium was replaced with StemPro-34 containing SCF (50 ng/mL), IL-3 (10 ng/mL), TPO (50 ng/mL), Flt-3 ligand (10 ng/mL), IL-6 (10 ng/mL), IL-11 (5 ng/mL), IGF-1 (25 ng/mL) and EPO (2 Units/mL) for LSEP induction. A diagram of differentiation protocol is included **Fig. 1a**. Venous angioblast (VA) induction was performed according to previously reported differentiation protocols until day 8^26^.

### Flow cytometry and cell sorting

Flow cytometric analyses and cell sorting were performed using the BD LSRFortessa^TM^ cell anlyzer (BD Bioscience) and FACSAria Fusion (BD Bioscience) or QuadroMACS Separator with LS Columns (Milteny Biotec) respectively. Antibodies and their dilution rates are as follows: anti-CD34 (BD Biosciences, 562577, 1:50), anti-CD32 (BioLegend, 303205, 1:50), anti-CD73 (Milteny Biotec, 130-095-183, 1:50), anti-CXCR4 (BioLegend, 306506, 1:50), anti-CD43 (BD Biosciences, 744662, 1:50), anti-CD31 (BD Biosciences, 740973, 1:50), anti-LYVE1 (R&D, AF2089, 1:50) conjugated to Alexa Fluor 647 by Lightning-Link (abcam), anti-EPCAM (BD Biosciences, 563180, 1:50), and anti-CD36 (BioLegend, 336207, 1:50). For CD34^+^ cell isolation, CD34 MicroBead Kit (Milteny Biotec, 130-046-702, 10 µl/10 million cells) was used. The data analyses were performed using the FlowJo software (Treestar).

### Liver Sinusoidal-like endothelial cell (LSEC) induction

For sinusoidal-like endothelium induction, day 8 adherent CD34^+^ iLSEP were dissociated and stained with CD34 MicroBead Kit (Milteny Biotec). After staining, cells were sorted using QuadroMACS Separator with LS Columns, then sorted cells were disseminated onto 24-well tissue plate coated with Matrigel Growth Factor reduced (Corning) at 1:50 dilution and cultured in a mixture of endothelial cell growth medium (EGM, Lonza) and hepatocyte culture media (HCM, Lonza) with Rock inhibitor (10 µM) and VEGF-A (5 ng/mL), and with or without OSM (10 ng/mL) at a density of 8.8e4 cells per well. After 24 hours culture, medium was replaced by same differentiation medium without Rock inhibitor. The cytokines were purchased from R&D systems or PeproTech. The cells were cultured under 37°C, with 5% CO2 and 5% O2. A diagram of differentiation protocol is included **Extended Data Fig. 2f**. LSEC-like cells from VA (VA- LSEC) were induced from VA according to our previously reported differentiation protocol from day 8 to day 12^26^. The cell seeding density on day 8 was changed to 8.8e4 cells per well for 24-well plates, in accordance with our LSEC induction protocol.

### Formation of HLBO by inverted multilayered air-liquid interface 3D (IMALI) culture

Directed differentiation methods for liver bud cells: hepatic endoderm (iHE), septum mesenchymal cells (iSTM), and arterial endothelial cells (iAEC) are described previously^24^.

The multilayer gradient gel culture system as previously described^45,46^ was modified as follows. Hanging cell culture inserts (polyethylene terephthalate membrane, pore size of 8.0 µm, Millipore) were placed upside-down in a sterile plastic box. For a multilayering gel technique, Matrigel diluted 1:1 with a mixture of EGM and HCM containing dexamethasone (0.1 mM; Sigma-Aldrich), oncostatin M (10 ng/mL), hepatocyte growth factor (HGF) (20 ng/mL, PromoKine), SingleQuots (Lonza), SCF (50 ng/mL), IL-3 (10 ng/mL), TPO (50 ng/mL), Flt-3 ligand (10 ng/mL), IL-6 (10 ng/mL), IL-11 (5 ng/mL), IGF-1 (25 ng/mL) and EPO (2 Units/mL) were sequentially applied to the bottom membrane surface of the insert. First, to construct the gel basement, a 5 µL empty gel drop was applied and the gel drop was solidified after 15 min incubation at 37 °C. Next, the first cell layer consisted of a 3 µL drop with day 10 liver bud cells at a final cellular concentration of 44 million cells/mL and a ratio of 10:7:1 (iHE / iAEC / iSTM) was placed on top of the previous drop. Gelling process was accomplished after 15 min incubation at 37 °C. Finally, the second cell layer consisted of an 8 µL drop with day 7 CD34^+^ purified iLSEP at a final cellular concentration of 16.5 million cells/mL was placed on top of the previous two drops. Gelling process was concluded after 20 min at 37 °C. Matrigel was always kept on ice to avoid undesired solidification. Inverted hanging cell culture insert with a multilayered gel-embedded cells were placed in Ultra-Low attachment 6 well plate and 2 mL of culture medium was added for each well. A diagram of procedures is included **Fig. 1a**. When identifying iLSEP and iAECs added in HLBO together, GFP reporter iPSC line and mCherry reporter iPSC line were used for fluorescent tagging, respectively.

### Immunofluorescent staining

Cells were fixed in 4% paraformaldehyde (Wako) in PBS for 15 min at 4 °C, then washed twice with PBS. Samples were permeabilized with 0.125% Triton-X (Sigma) for 45 min, and then blocked with Donkey serum (Millipore) for 30 min at room temperature followed by overnight incubation at 4 °C with primary antibodies in staining buffer (0.1% donkey serum in PBS). Primary antibodies and their dilution rates are as follows: anti-CD34 (Abcam, ab81289, 1:50), anti-CD31 (Abcam, ab24590, 1:100), anti-CD32b (Abcam, ab151497, 1:500) and anti-FVIII (Abcam, ab41188, 1:100). Next day, samples were washed twice with wash buffer (0.1% tween 20 (Sigma) in PBS), incubated with species-specific secondary antibodies conjugated with Alexa Fluor 555, 594, or 647 (Life Technologies) and DAPI (Sigma) for nuclear staining diluted 1:1000 in staining buffer for one hour at room temperature, and then washed again twice with wash buffer. Images were acquired using KEYENCE BZ-X fluorescent microscope (KEYENCE) or LSM 880 with Airy scan (Zeiss). 2D images processing was managed by Image J software (NIH) with Fiji software package. Cellpose 2.0 software (https://github.com/mouseland/cellpose)^80^ was used for CD31^+^ and/or CD32b^+^ endothelial cell segmentation and counting.

### Scanning electron microscopy of endothelial cells

Day 12 iLSEC and day 10 iAEC treated with OSM were fixed with a mixture of 4% paraformaldehyde and 0.5% glutaraldehyde (Wako) in 0.1 M phosphate-buffer (PB) for 2 hours at 4 °C and then quenched by 50 mM NH4Cl/PBS for 15 min. They were washed overnight at 4 °C in the same buffer and post-fixed with 1% OsO4 buffered with 0.1 M phosphate-buffer for 2 hours. The specimens were dehydrated in a graded series of ethanol and dried in a critical point drying apparatus (JCPD-5; JEOL) with liquid CO2. They were spatter-coated with platinum and examined by scanning electron microscopy (JSM-7900F; JEOL, Tokyo, JAPAN).

### scRNA-seq library preparation and sequencing

At days 3, 7, and 11 after self-assembly culture, three HLBOs at each time point were subjected to scRNA-seq library preparation. To dissociate into single cells, HLBOs attached to transwell membranes were harvested with a cell scraper and transferred to 2 mL tubes. The HLBOs were washed three times with HBSS without Ca2+ and Mg2+, then 500 μL of a dissociation solution containing 0.1 mg/mL Liberase TM (Roche) and 0.1 mg/mL DNase I (Sigma) diluted in DMEM/F12 medium was added. The mixture was incubated at 37°C for 30 minutes with shaking at 100 rpm. The HLBOs were physically dissociated by pipetting until no cell clumps were visible. The dissociated cells were then washed with 500 μL PBS containing 1% BSA and centrifuged at 300g for 5 minutes. The cell suspension was filtered through a 70 μm strainer, centrifuged again, and resuspended in PBS containing 1% BSA. Cell counting was performed, and targeting 10,000 cells, the cell suspension was aliquoted and subjected to single-cell droplet formation using the 10x Chromium platform. The library preparation was performed using the NextGEM Single Cell 3’ v3.1 kit (10x Genomics) according to the manufacturer’s instructions. Sequencing was performed on the Illumina NovaSeq 2x150 bp configuration for 1 lane (with 1%Phix).

### scRNA-seq raw data processing

The raw scRNA-seq demultiplexes raw base call (BCL) files were converted to FASTQ files by the bcl2fastq function, and the count pipeline implemented in Cell Ranger Software Suite version 2.0 (10x Genomics). was used to align and quantify the sequencing reads, obtaining feature-barcode and raw UMI count matrix. All UMI count matrices were subjected to primary data processing using the Seurat v5 package implemented in R. Cells were filtered based on the following exclusion criteria: unique feature counts less than 500 or total RNA count less than 2000, mitochondrial count greater than 20%, ribosomal count greater than 40% and less than 5%. Genes which were expressed in fewer than 3 cells were also removed. DoubletFinder was applied to remove any doublet cells. The data was normalized using default global-scaling method. The cell cycle phase was estimated using the “CellCycleScoring()” function. All HLBO data were aggregated into a single object for integration using the “merge()” function and then output. The output data was reloaded in Scanpy to perform subsequent analysis in the Python environment. Data integration was performed using scPoli^81^. In the scPoli model, we changed the following parameters from the default configuration for integration: embedding_dim was set to 30; hidden_layer_sizes were determined as the square root of the total number of cells. During the training phase, we employed the default configuration. Three hierarchical levels of annotation were assigned to the identified clusters: cell class (Layer 1), which is the upper domain grouping cell types; cell type (Layer 2); and subtype (Layer 3). These annotations were guided by known cell type markers in primary liver and the automatic annotation results using the decoupleR package implemented in Scanpy.

### Constructing an integrated dataset of published human liver organoids and a primary liver reference

A publicly available, 11 different protocol-based human PSC-derived liver organoid datasets were collected according to the descriptions in the original publication^2,13,53–58^ (**Supplementary Table 6**). Briefly, we obtained available data (either raw FASTQ files, count matrices, or Cell Ranger outputs such as “filtered_feature_bc_matrix” files) for each organoid from databases such as GEO, ArrayExpress, and the Human Cell Atlas (HCA). For primary adult and fetal datasets^82^, we downloaded H5AD data deposited in the HCA (https://collections.cellatlas.io/liver-development). For FASTQ files, we used Cell Ranger to align and quantify the sequencing reads with the same parameters described in the original publication, generating UMI count data. Subsequent data processing was performed in Scanpy using default settings. We curated metadata for all organoid data, including cell barcodes, sample names, cell type annotations, and cell cycle phase. We normalized and combined the public organoid data with our HLBO data. cell type annotations were based on the original publication and assigned into hepatocytes, hepatoblasts, endothelial cells, cholangiocytes, and stellate cells, which were added as new metadata. Concurrently, we identified the top 3,000 variable genes from the primary liver data and applied these to the organoid dataset. Using scPoli, we integrated the organoid data comprising 91,700 cells and the primary liver data comprising 16,187 cells. The same scPoli configulations were used as integrating the HLBO data. After integration, we performed Louvain clustering and re-annotated cell types based on the expression of known marker genes. For further comparative analyses, we used the integrated organoid data as the query and the primary liver data as the reference.

### Comparative analysis with primary adult and fetal liver

To benchmark our HLBO model against existing models, we used miloR^60^ and scrabbitr^59^ R package to compute neighborhood graph correlation mapping and spectrum comparison for primary adult and fetal liver dataset. The neighborhood correlations were computed using 3000 highly variable genes that were found in the highly variable genes in either adult or fetal primary liver compared as reference. The transcriptional similarity graph was computed using 30- dimensional nearest neighbors and UMAP embeddings of cells. Other parameters were implemented as default.

### RNA velocity analysis

BAM files aligned using the count pipeline implemented in Cell Ranger were first sorted by using SAMtools v1.6. Next, the Velocyto pipeline was used to count spliced and un-spliced reads and generate loom files of each HLBO samples. Gene-specific velocities were then computed following the scVelo package^83^. The velocity streams predict by scVelo with the loom files were embedded in UMAPs that plotted endothelial cells colored by subtype. Additional developmental trajectory inference algorithm, partition-based graph abstraction (PAGA), was used. Kinetic plots for the latent time and the expression of various individual gene were generated based on the velocity calculated by scVelo.

### Gene set enrichment analysis

From the integrated scRNA-seq data of organoids, hepatocytes were extracted and differentially expressed genes (DEGs) between HLBOs in this study and publically available organoids were identified by “scanpy.tl.rank_genes_groups()” function in scanpy (pvals_adj < 0.05). GSEA using the DEGs was performed by GSEApy. Enrichment plots were generated using the function implemented in GSEApy.

### RNA isolation and Bulk RNA-seq

Total RNA was extracted from cultured cells, using a PureLink RNA Mini Kit (Thermo Fisher Scientific). A cDNA library was generated from 10 ng of total RNA, using a SuperScript VILO cDNA Synthesis Kit (Thermo Fisher Scientific).

Amplification, primer digestion, and adapter ligation were performed using an Ion AmpliSeq Transcriptome Human Gene Expression Kit (Thermo Fisher Scientific). The cDNA library was purified using Agencourt AMPure XP Reagent (Beckman Coulter) and quantified using Ion Library TaqMan Quantitation Kit (Thermo Fisher Scientific), followed by dilution to 75 pM with water and pooled equally. Eight samples per pool were sequenced using an Ion 540 Chip Kit (Thermo Fisher Scientific) simultaneously, using IonS5 XL (Life technologies) and Ion Chef instrument systems (Life technologies) with Ion 540 Kit-Chef (Life technologies). All procedures were performed according to the manufacturers’ protocols.

### Bulk RNA-seq raw data processing

For the primary human liver tissue and one cryopreserved primary human LSEC isolated from tissue, raw data obtained with the same RNA isolation and sequencing methods and deposited in GSE152447 of GEO were used for subsequent data processing and analysis. From the sequencing output, quality control, alignment reads on hg19 (hg19_AmpliSeq_Transcriptome_21K_v1.bed), read counts and the normalization (reads per million, RPM) for each gene were obtained using the Torrent ampliSeqRNA plugin (hg19 AmpliSeq Transcriptome ERCC v1) in Torrent Suite Software v5.2.1. Data scaling of AmpliSeq count data were performed by converting into z-score.

### Functional classification, clustering and visualization of bulk RNA-seq data

Scaled dataset were first subjected to gene functional classification based on gene sets related to cytokines, growth factors and drug metabolic process. The list of gene sets derived from the Gene Ontology (GO) (Gene Ontology Consortium) of Molecular Signatures Database (MSigDB, v7.0). Unsupervised clustering using Euclidean distance and average linkage method and visualization of heatmap and dendrogram were performed by R package heatmap.2.

### Intercellular communication analysis

We applied scaled dataset of bulk RNA-seq or scRNA-seq as Seurat object to CellChat R package^84^. The annotation of ligand-receptor pair was automatically assigned based on “Secreted Signaling” subset of CellChatDB.human database. The probability of intercellular communication and significantly enriched ligand-receptor pairs were inferred by the “timean” method with default parameters. Visualization was done using the functions implemented in CellChat.

### qRT-PCR analysis

Reverse transcription was carried out using the High-Capacity cDNA Reverse Transcription Kit (Applied Biosystems) according to the manufacturer’s protocols. qPCR was carried out using TaqMan gene expression master mix (Applied Biosystmes) on a QuantStudio 3 Real-Time PCR System (Thermo Fisher Scientific). All primers and probes information for each target gene were obtained from Custom TaqMan Probes (https://www.thermofisher.com/order/custom-oligo/custom-taqman-probes).

### Whole mount clearing and imaging

Organoids were fixed in 4% paraformaldehyde in PBS for 2 hours at 4 °C. Clearing and immunofluorescent staining were performed by using SCALEVIEW-S (Wako) according to AbSca*l*e protocol supplied from manufacturer. Primary antibodies and their dilution rates are as follows: anti-CD31 (1:50), anti-CD32b (1:500), anti-FVIII (1:100), anti-ASGR1 (R&D, MAB4394, 1:100), anti-GS (Abcam, ab73593, 1:500), anti-LYVE1 (R&D, AF2089, 1:100), anti-CYP3A4 (Thermo Fisher Scientific, MA5-17064, 1:500), anti-PXR (Thermo Fisher Scientific, PA5-72551, 1:25), and anti-CYP1A2 (Abcam, ab22717, 1:100). Species-specific secondary antibodies conjugated with Alexa Fluor 488, 555, or 647 (Life Technologies) were used at 1:500 dilution. DAPI (Sigma) for nuclear staining was diluted 1:1000. z-stack images were acquired using LSM 880 with Airy scan (Zeiss). 3-D image processing and volume rendering were managed by Imaris software (Oxford Instruments). Quantification of fluorescence signal for z-stack images was performed using Image J software (NIH) with Fiji software package.

### Fetal samples

All human fetal samples were collected under Institutional Review Board Approval (STUDY-22-00065) at the Icahn School of Medicine at Mount Sinai. Consent for tissue donation was obtained after the patient had already made the decision for pregnancy termination and was obtained by a different clinical research coordinator than the physician performing the procedure. All tissues were de-identified, and the only clinical information collected was gestational age and the presence of any maternal or fetal diagnoses.

### H&E staining and immunohistochemistry

Organoids were fixed in 4% paraformaldehyde in PBS for 2 hours at 4 °C and tissues were immersed in 10% neutral buffered formalin for fixation. Fixed organoids and tissues were frozen at -80°C, embedded in Tissue-Tek optimal cutting temperature (OCT) compound (SAKURA), and cut to 7-10 μm thick sections on glass slides. Hematoxylin and eosin (H&E) staining was performed using standard protocols.

Immunohistochemistry was performed using the same procedure as for immunofluorescence staining. Primary antibodies and their dilution rates are as follows: anti- THBD (Invitrogen, MA5-11454, 1:100), anti-LYVE1 (R&D Systems, AF2089, 1:100), anti-vWF (Abcam, ab179451, 1:500), anti-GS (GeneTex, GTX630657, 1:250), anti-GLS2 (Abcam, ab113509, 1:100), anti-STAB1 (Novus Biologicals, H00023166-M05, 1:50), anti- CD32b (Abcam, ab151497, 1:500), anti-Ku80 (CST, #2180, 1:400), anti-human nuclear antigen (Novus NBP2-34342, 1:200) and anti-human-specific Albumin (Sigma, A3293, 1:200). Species-specific secondary antibodies conjugated with Alexa Fluor 555, 594, or 647 (Life Technologies) and DAPI (Sigma) for nuclear staining was diluted 1:1000. Images were acquired using KEYENCE BZ-X microscope (KEYENCE) or LSM 880 with Airy scan (Zeiss), and fluorescence signals were processed and quantified by Image J software (NIH) with Fiji software package. CellProfiler (Broad Institute) was used for segmentation and counting of ASGR1^+^ and/or GS^+^ and/or GLS2^+^ hepatocyte normalized by counterstaining (DAPI).

### Transmission immunoelectron microscopy of HLBO

HLBOs were fixed with 4% paraformaldehyde in PBS overnight at 4 °C. The fixed organoids were embedded in OCT compound at -80°C, and cut to 7 μm thick sections on glass slides. The cryosections were washed with 0.1 M PB, permeabilized with block solution (50 mM NH4Cl, 0.1% saponin, 1% bovine serum albumin in 0.1 M PB), and incubated with anti-LYVE1 antibody (ab36993; abcam; 1:100 dilution) and goat anti-rabbit IgG Fab’ fragment antibody conjugated with 1.4 nm Nanogold particles (#2003; Nanoprobes). The pre-embedding silver enhancement immunogold method was used as previously described^85^. After the labelling, the samples were washed with 0.1 M PB three times, postfixed in 1% osmium tetroxide in 0.1 M PB for 2 hours, dehydrated, and embedded in Epon 812 according to a standard procedure. Ultrathin sections were stained with uranyl acetate and lead citrate and observed using a JEM- 1400Flash electron microscope (JEOL).

### siRNA transfection

The mixture of Silencer™ Select Wnt2 siRNA (s14865, s14866, s14867) was used for knockdown of wnt2, and Silencer™ Select Negative Control No. 1 siRNA was used as a negative control in parallel. For fluorescent tracking of siRNA, BLOCK- iT™ Alexa Fluor™ Red Fluorescent Control (Thermo Fisher Scientific) was used. siRNA was formulated into lipid-based nanoparticles (LNPs). In the gelling process for IMALI culture, LNPs-siRNA complex was mixed with CD34^+^ purified iLSEP in gel at a final concentration of 200 nM.

### Albumin, complements, VEGF-A and FVIII Enzyme-linked immunosorbent assay (ELISA)

For measurement of Human albumin (ALB), complements, VEGF-A and FVIII, 900 μL of culture supernatant of organoids on Ultra-Low attachment 6 well plates (Corning) were collected and stored at - 80°C until use. The supernatant was assayed with a Human Albumin ELISA Quantitation Kit (Bethyl Laboratories Inc.), Human C3, C5, Factor H and Factor D ELISA Kit (abcam), LBIS Human VEGF ELISA Kit (FUJIFILM), and Human Coagulation Factor VIII (F8) ELISA Kit (Biomatik Corporation) according to the manufacturer’s protocols.

### Analysis of glutamine metabolism

300 μL of HLBO lysates were prepared by T-PER Tissue Protein Extraction Reagent (Thermo Fisher Scientific) with protease inhibitor (xxx). The amount of protein in the lysate was quantified by NanoOrange™ Protein Quantitation Kit (Thermo Fisher Scientific) and the enzymatic activity of glutamine synthetase was measured by Glutamine Synthetase Activity Assay Kit (Abcam). The supernatant of HLBO was assayed with Glutamine/Glutamate-Glo Assay (Promega) and Ammonia Assay Kit (Abcam). Ammonia uptake in 24 hours was quantified by the difference between the HLBO supernatant and the culture medium.

### Analysis of activated partial thromboplastin time (APTT)

30 μL of culture supernatant of organoids, rhFVIII, or medium for a negative control were mixed with 270 μL of CRYOcheck factor (FV, VIII, IX and XI) deficient plasma (Precision BioLogic). For a positive control, 30 μL of medium was mixed with 270 μL of CRYOcheck Normal Reference Plasma. For testing APTT, they were subjected to the CA-620 automated hemostasis analyzer (Sysmex) according to the manufacturers’ protocols. Comparisons between samples were made using actual values (sec) or the percentage shortening of the APTT relative to the negative control with the positive control as the maximum value. Negative and positive controls were obtained for every measurement and were used for normalization to the data at the same time of measurement.

### Fibrin clot

5 mg/mL Fibrinogen (BioVision) and 5 mg/ml Alexa Fluor 488-conjugated fibrinogen (invitrogen) diluted in 50 mM Tris/100 mM NaCl were mixed in 25 μL each on 96- well plates and incubated at 37°C for 48 hours. After incubation, 25 μL of 5 mM CaCl2 buffer, 1 μL of organoid culture supernatant, and 25 μL of CRYOcheck factor VIII deficient plasma (Precision BioLogic) were added to each well. The mixture was immediately placed on each glass slide, cover-slipped and incubated at room temperature for 2 hours. Images of fibrin clot were acquired using LSM 880 with Airy scan (Zeiss). As a positive control, rhFVIII was used instead of culture supernatant.

### Animals

All animal experiments, bred, and maintaining were conducted in accordance with the ethical guidelines and protocol approved by the Institutional Animal Care and Use Committee of Takeda Pharmaceutical Company Limited (AU-00031097 and AU-00030165) and Tokyo Medical and Dental University (A2022-133C). All animals were housed individually under controlled temperature (20–26 °C), humidity (40–70%), and a 12 h light-dark cycle (lights on 7:00–19:00). Female non-obese diabetic/severe combined immunodeficient (NOD/SCID) mice for intravital imaging under a cranial window were purchased from CLEA Japan, Inc.. Male NOD/Shi-scid,IL-2RγKO Jic (NOG) mice for pharmacokinetic analysis after HLBO transplantation were purchased from In-Vivo Science Inc.. For Hemophilia A *in vivo* efficacy and transplantation study, we generated immunodeficient NS-IL2rg KO/ FVIII KO Hemi (NSI-FVIII KO) mice. First, we generated NOD/scid/*Il2rg*^-/-^ (NSI) mice by genome editing technology using Zinc finger nuclease (ZFN). Pre-validated CompoZr(R) ZFN plasmids for *Il2rg* (CKOZFN33739) were purchased from Sigma-Aldrich. Il2rg ZFN mRNAs were synthesized by *in vitro* transcription with mMESSAGE mMACHINE T7 Ultra kit (Thermo Fisher Scientific). The mRNAs were injected into the pronuclei of zygotes which were obtained by standard *in vitro* fertilization method (IVF) using sperm and eggs of NOD/scid [NOD.CB17-Prkdc-scid/J] mice (Charles River Laboratories Japan). A female heterozygous *Il2rg* knock-out mouse was mated with a male NOD/scid mouse to generate the NSI background mice. Next, we knocked out the factor VIII gene (*F8*) from the NSI mice using the CRISPR/Cas9 system. A diagram of *F8* gene editing is included **Extended Data Fig. 10f**. For details, the Cas9 coding sequence was added downstream of the T7 promoter sequence by PCR. The amplified PCR fragment was ligated into pMD20-T vector (Takara Bio). The Cas9 plasmid was used as a template for *in vitro* transcription using mMESSAGE mMACHINE T7 Ultra Kit. Two single-guide RNA (sgRNA) sequences were designed to sandwich the exons 17 and 18 of *F8*. T7 promoter was added to each of the sgRNA sequences by PCR. The amplified PCR fragments were used as templates for *in vitro* transcription using MEGAshortscript T7 Kit (Life technologies). The single-stranded oligodeoxynucleotide (ssODN) containing homologies of 60 bases on both sides flanking each of the sgRNA targeting sequences was purchased as Standard Oligo (Eurofins Genomics). For microinjection, NSI background zygotes were obtained from NSI mice by IVF. A mixture of Cas9 mRNA (100 ng/ul), two sgRNAs (50 ng/uL) and ssODN (50 ng/uL) was injected into the pronuclei of zygotes. An obtained male *F8* knock-out mouse was mated with female NSI mice to generate the female NSI-F8^-/+^ mice.

### Intravital imaging of HLBOs under a cranial window

The cranial window was prepared as described previously. 8- to 14-week-old mice were used for preparing the cranial window. Three HLBOs cultured for 3 days after assembly *in vitro* were collected and implanted on the mouse cortex under a cranial window of 9- to 16-week-old mice as described previously.

Intravital imaging was performed using an Olympus FV3000 confocal laser scanning microscope or a Nikon AX R confocal microscope with high-speed resonant scanning system. The perfused vessels were highlighted by tail vein injection of MW 2,000,000 tetramethyl-rhodamine-labeled dextran (Thermo Fisher Scientific, D7139) or MW 70,000 Texas Red-labeled dextran (Thermo Fisher Scientific, D1864). Time-lapse imaging of 70 kDa dextran permeability was performed starting at 3 minutes after injection, with images acquired every 3 minutes for 60 minutes. For staining, Alexa Fluor 647-conjugated anti-mouse-specific CD31 antibody (BioLegend, 102516) or AcLDL (Thermo Fisher Scientific, L35354) conjugated to Alexa Fluor 647 by Lightning-Link (Abcam, ab269823) were tail vein injected. The excitation wavelength or laser power must be adjusted to each fluorescent sample at every experimental setting.

### Transplantation of HLBOs into liver

NSI-FVIII KO mice were anesthetized inhaled isoflurane and placed on temperature-controlled heating pads (37 °C). To temporarily rescue bleeding during operations due to FVIII deficiency, 200 µL of 50 Units/kg rhFVIII was administered intravenously. For transplantation into liver, the implantation site was prepared by shaving about 1 cm of the liver surface, and after hemostasis with cotton, three organoids cultured for 3 days after assembly *in vitro* were placed and immediately glued using tissue adhesive (Beriplast P Combi-Set, CSL Behring). The abdominal wall and the skin incision were closed with suture. Immediately after closure, all mice received a single injection of meloxicam (METACAM, Boehringer Ingelheim) at 5 mg/kg for pain management. Post-implantation mice were followed for up to 20 weeks, at which point they were euthanized.^24^

### Bleeding assay

Mice were anesthetized with inhaled isoflurane (Pfizer) and placed on temperature-controlled heating pads (37 °C), and the distal portion of the tail was cut at 1.5- mm diameter (moderate hemostatic challenge), after which the tail was immersed in a predefined volume of 37 °C saline (0.9% NaCl) for 25 min. Amount of blood loss was determined by the hemoglobin concentration in the saline solution after red cell lysis with 2% acetic acid and measured by Absorbance at 405 nm. Groups of NSI-FVIII KO mice were injected intravenously (retro-orbital) within 3 min before the tail cut with equal volumes (200 μL) of dialyzed supernatant of organoids which was ultra-filtered into approximately 12-24- fold concentration using AMICON ULTRA-15 10KDA ULTRACEL-PL MEMBRANE (Merck), 50 Units/kg rhFVIII, or saline (Sham) diluted in sterile sodium chloride 0.9% for injection.

## QUANTIFICATION AND STATISTICAL ANALYSIS

Statistical analysis was performed using GraphPad Prism 7. Data were statistically evaluated from biological independent replicates. The details of statistics and exact number of replicate (*n*) are shown in Figure legend. Briefly, for two-samples comparison, unpaired two-tailed Welch’s t-test, Mann-Whitney’s U-test, Wilcoxon’s rank-sum test was used. If repeated tests were performed, p-values were adjusted by Bonferroni or Holm-Sidak method or cutoff by the false discovery rate (FDR) < 10%. For comparisons between more than two samples, one-way or two-way analysis of variance (ANOVA) with Dunnett’s or Tukey’s, or Holm-Sidak’s multiple comparisons test. p-values or adjusted p-values < 0.05 were considered statistically significant.

## Data availability

RNA-seq raw and processed data have been deposited in the Gene Expression Omnibus (GEO) under accession numbers GSE217649 (Bulk) and GSE270807 (single cell) respectively. The data that support the findings of this study are available from the corresponding author upon reasonable request.

## ACKNOWLEDGMENTS

The authors would like to express their sincere gratitude to S. Yamanaka and S. Izumo, Y. Kajii, and A. Nakanishi for supporting the collaboration research between Takeda Pharmaceutical Company and Center for iPS Cell Research and Application (CiRA), Kyoto University (T-CiRA). We thank T. Shinozawa and H. Anayama (Takeda Pharmaceutical Company) for advice on the research strategy; Y. Noguchi, M. Fukui, A. Nukuda (Orizuru Therapeutics, Inc.) for technical assistance with *in vitro* experiments; M. Ide and M. Nomura (RABICS, LTD.) for technical assistance with animal experiments. We also thank K. Hashikami, T. Yamamura, and M. Takeyama (Axcelead Drug Discovery Partners, Inc.) for generating and providing NSI-FVIII KO mice; T. Sasaki and K. Aoyama (Axcelead Drug Discovery Partners) for LC-MS/MS analysis; C. Moriya (TMDU) for providing NOD-SCID mice with cranial window; Y. Sakamaki (TMDU) for her expert technical assistance and advice with electron microscopy; A. Niwa (CiRA) for advice and discussion on differentiation of the hemogenic endothelium; H. Matsumoto (T-CiRA), M. Maezawa, and M. Mori (TMDU) for kind administrative assistance. This work was supported by the T-CiRA Joint Program from Takeda Pharmaceutical Company. T. T. is a New York Stem Cell Foundation – Robertson Investigator. This work was also supported by Cincinnati Children’s Research Foundation grant, CURE award, NIH Director’s New Innovator Award (DP2 DK128799-01), NIH UG3/UH3 DK119982, PHS Grant P30 DK078392 (Integrative Morphology Core and Pluripotent Stem Cell and Organoid Core) of the Digestive Disease Research Core Center in Cincinnati, the Falk Transformational Awards Program, JST Moonshot R&D Grant Number JPMJMS2033 and JPMJMS2022, Takeda Science Foundation award, Mitsubishi Foundation award and AMED CREST (JP22gm1210012), JP20fk0210037, JP20bm0704025, JP21fk0210060, JP21bm0404045, JP22fk0210091, JP22fk0210106, JST JPMJPS2033, JPJSBP220203101, and JSPS JP18H02800. Fetal tissue analysis was conducted with the support of the Developmental Origins of Health and Disease Biorepository at Icahn School of Medicine at Mount Sinai. We are grateful to the Biorepository laboratory team and Biorepository participants for their contributions to this research.

## AUTHOR CONTRIBUTIONS

N.S., and T.T. conceived the study. N.S., Y.N., and T.T. designed the experiments; N.S. wrote original draft of the manuscript; Y.Y., T.L.W., and T.T. reviewed and edited the manuscript; N.S. performed almost all *in vitro* and animal experiments, bioinformatics, and analyzed data; Y.N. performed and optimized hemophilia A animal model experiments; Y.Y. performed scanning electron microscopy experiment; S.K., performed cranial window animal experiments; E.K. and A.K. performed and optimized fibrin clot experiment; J.F. and T.S. performed the cell culture and biochemical assays; K.I., R.O., P. C-E., and Y-W. C. performed sectioning, staining, and analysis; T.K., M.K. obtained and analyzed some RNA-seq data; M.F. performed and optimized siRNA transfection experiment.

